# In vitro reconstitution of divisome activation

**DOI:** 10.1101/2021.11.08.467681

**Authors:** Philipp Radler, Natalia Baranova, Paulo Caldas, Christoph Sommer, Mar López-Pelegrín, David Michalik, Martin Loose

## Abstract

Bacterial cell division is coordinated by the Z-ring, a cytoskeletal structure of treadmilling filaments of FtsZ and their membrane anchors, FtsA and ZipA. For divisome maturation and initiation of constriction, the widely conserved actin-homolog FtsA plays a central role, as it links downstream cell division proteins in the membrane to the Z-ring in the cytoplasm. According to the current model, FtsA initiates cell constriction by switching from an inactive polymeric conformation to an active monomeric form, which then stabilizes the Z-ring and recruits downstream proteins such as FtsN. However, direct biochemical evidence for this mechanism is missing so far. Here, we used biochemical reconstitution experiments in combination with quantitative fluorescence microscopy to study the mechanism of divisome activation *in vitro*. By comparing the properties of wildtype FtsA and FtsA R286W, a gain-of-function mutant thought to mimic its active state, we found that active FtsA outperforms the wildtype protein in replicating FtsZ treadmilling dynamics, filament stabilization and FtsN recruitment. We could attribute these differences to a faster membrane exchange of FtsA R286W as well as its higher packing density below FtsZ filaments. Using FRET microscopy, we also show that binding of FtsN does not compete with, but promotes FtsA self-interaction. Together, our findings shed new light on the assembly and activation of the bacterial cell division machinery and the mechanism of how FtsA initiates cell constriction.

## Introduction

Bacteria have intricate intracellular organizations, where different proteins localize to distinct sites in a tightly regulated, highly dynamic manner. The molecular mechanisms that give rise to these complex spatiotemporal dynamics are often unknown. This is in particular true for the divisome, a highly complex protein machinery that accomplishes cell division with remarkable precision^1^. The divisome consists of more than a dozen different proteins that assemble in a step-like manner. Divisome assembly is initiated by the simultaneous accumulation of FtsZ, FtsA and ZipA at midcell, where they organize into the Z-ring, a composite cytoskeletal structure of treadmilling filaments at the inner face of the cytoplasmic membrane **(Fig. 1a)**. In a second step, this dynamic Z-ring recruits cell division proteins to the division plane and promotes their homogeneous distribution around the circumference of the cell^2^. Finally, the cell starts to constrict while generating two new cell poles splitting the dividing cell in two. Although the biochemical network underlying cell division is now well studied^3^, how the membrane anchors of FtsZ control the timing of recruitment and activation of cell division proteins located in the cell membrane is currently unknown^4–6^.

**Figure 1:**
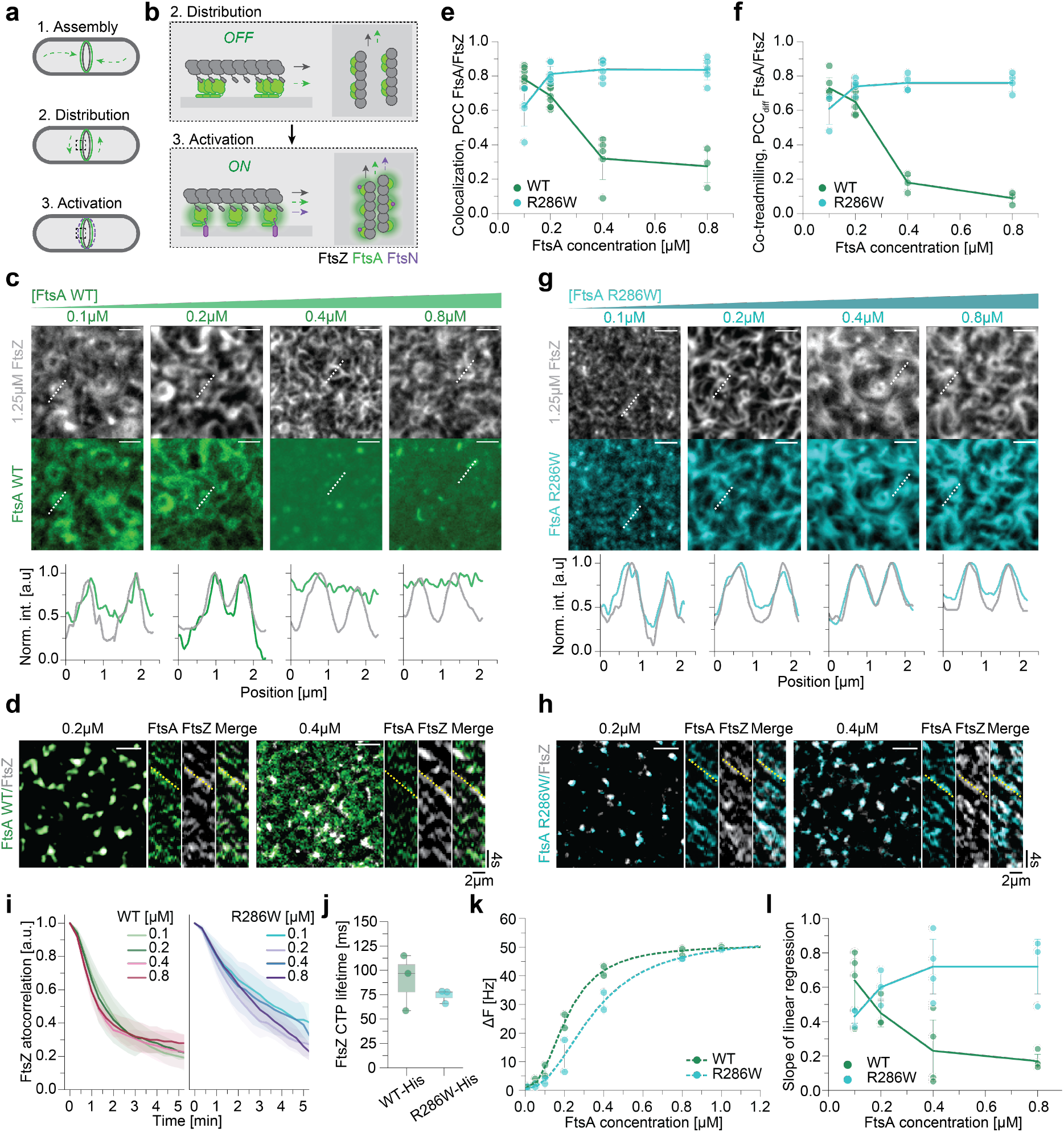
Membrane patterning of FtsA by treadmilling filaments of FtsZ. **a, b**, FtsA anchors FtsZ filaments to the cytosolic membrane of *E. coli*. Treadmilling of FtsZ distributes FtsA around midcell. **b**, Binding of FtsN is thought to switch FtsA from an oligomeric *off* state to the monomeric *on* state. FtsA activation triggers recruitment of divisome proteins and constriction. **c**, Representative micrographs of A488-FtsZ (grey) and Cy5-FtsA WT (green) at increasing FtsA WT and constant FtsZ concentration (1.25 μM). Intensity profiles correspond to dashed white lines. **d**, Representative micrographs showing merged differential images of FtsA WT with FtsZ for 0.2 μM (left) and 0.4 μM (right) FtsA. Yellow lines in kymographs indicate the slope for treadmilling FtsZ. Above 0.2 μM FtsA WT fails to replicate FtsZ dynamics. **e**, Colocalization of FtsA/FtsZ quantified by PCC shows that colocalization of FtsA WT (green) and R286W (cyan) with FtsZ starts to differ significantly at 0.4 μM (0.32 ± 0.12; 0.84 ± 0.05; p-value: 1.28*10^−5^). **f**, Dynamic colocalization of FtsA/FtsZ quantified by PCC_diff_. FtsA R286W follows FtsZ treadmilling more efficiently than FtsA WT at FtsA concentrations above 0.2 μM (0.18 ± 0.04; 0.76 ± 0.04; p-value: 1.82*10^−4^). **g**, Representative micrographs of A488-FtsZ (grey) and Cy5-FtsA R286W (cyan) at increasing FtsA R286W and constant FtsZ concentration (1.25 μM). FtsA R286W colocalizes with FtsZ filaments at all tested concentrations. Intensity profiles correspond to dashed white lines. **h**, Representative micrographs showing merged differential images of FtsA R286W with FtsZ for 0.2 μM (left) and 0.4 μM (right) FtsA. Yellow lines in kymographs indicate the slope for treadmilling FtsZ. FtsA R286W replicate FtsZ dynamics more robustly. **i**, The FtsZ network is more persistent in the presence of FtsA R286W, indicated by a slower decay of the autocorrelation. FtsZ persistency decreases slightly with higher FtsA WT concentrations (left), whereas it remains stable with FtsA R286W (right). **j**, The FtsZ-C-terminal peptide has the same lifetime on FtsA WT-His6 or R286W-His6 on membranes with 1% Tris-NTA lipids (90.16 ± 23.53ms; 74.01 ± 5.80ms; p-value: 0.40). **k**, QCM-D experiments reveal that FtsA R286W binds slightly weaker to bilayers compared to FtsA WT (0.32 ± 0.05μM vs 0.21 ± 0.01μM, p-value: 0.15). However, the membrane density is saturated with both FtsA variants at 0.8 μM. **l**, The slope of the linear regression is proportional to the [FtsZ] vs. [FtsA] ratio. With increasing FtsA WT concentrations this slope decreases, while it remains high for FtsA R286W. Scale bars in all micrographs are 2μm.

The actin-homolog FtsA is widely conserved and generally considered to be the more important membrane linker for FtsZ filaments^7,8^. It can reversibly bind to the membrane via a C-terminal amphipathic helix, where it recruits FtsZ filaments by binding to their C-terminal peptides **(Fig. 1b)**. FtsA was found to self-interact to oligomerize into actin-like single or double protofilaments^9^ as well as membrane-bound minirings composed of 12 FtsA monomers with a diameter of about 150 nm^10^. In addition, FtsA binds to many other proteins of the divisome, including FtsN, FtsQ, FtsX and FtsW^4,6,11–15^ highlighting its indispensable role for cell division. While FtsA is essential in *E. coli*, several FtsA mutants have been identified that can compensate for the loss of other essential cell division proteins, including the alternative membrane anchor ZipA, but also FtsEX, FtsN, FtsQ and FtsK^7,16,17^. *In vivo*, these mutants facilitate the recruitment of division proteins and stabilize the Z-ring, which can lead to premature division^5,8,16–18^. Importantly, these properties correlate with a reduced self-interaction in yeast two-hybrid assays as well as the absence of cytoplasmic rods when membrane-binding deficient proteins are overexpressed^17^. As suppressor mutations are located at or near the binding interface between two FtsA subunits, these observations led to a model, where FtsA oligomerization and recruitment of downstream proteins are mutually exclusive and where divisome maturation and cell constriction depends on the switch of FtsA from an inactive, polymeric state to the active, monomeric form^3,14,17^ **(Fig. 1b)**. While this model is consistent with many observations made *in vivo*, direct biochemical evidence for FtsA’s different activity states, their molecular properties and the mechanism of their conversion remains missing so far.

Here, we have reconstituted the dynamic interactions between treadmilling filaments of FtsZ, its membrane-anchor FtsA and the cytoplasmic peptide of the late division protein FtsN on membrane surfaces *in vitro*. By comparing the properties of wildtype FtsA and FtsA R286W, a hyperactive mutant that represents the activated state of the protein, we provide answers to two fundamental questions about the role of FtsA for divisome maturation and initiation of cell constriction: first, what is the relationship between the activity state of FtsA, its self-interaction and the recruitment of downstream proteins? And second, how does activation of FtsA affect the spatiotemporal organization of itself and that of FtsZ filaments on the membrane? By answering these questions, we shed light on the mechanism of bacterial cell division and also identify general requirements we believe to be important for the propagation of biochemical signals in living cells.

## Results

### Membrane patterning of FtsA by treadmilling filaments of FtsZ

FtsA localization to the division septum during the cell cycle is known to be FtsZ-dependent^19^. FtsZ also determines the circumferential dynamics of FtsA during treadmilling^20^. To study how FtsZ filaments direct binding of FtsA to the membrane on these two different time scales, we used a previously established *in vitro* reconstitution assay^4,21^ based on dual-colour TIRF imaging of proteins binding to a glass supported lipid bilayer. Using this approach, we were able to simultaneously record the dynamics of fluorescently labelled FtsZ and FtsA on the membrane surface at high spatiotemporal resolution.

When we added fluorescently labelled FtsZ and FtsA to the supported membrane at a concentration ratio similar to the one found *in vivo*^22^ (5:1) and lower (i.e. FtsZ = 1.25 μM and FtsA = 0.2 μM or 0.1 μM, with 75% A488-FtsZ and 66% Cy5-FtsA) the proteins immediately formed a dynamic cytoskeleton pattern of treadmilling filaments where both proteins closely overlapped **(Fig. 1c)**. We quantified the colocalization of the two fluorescent signals at steady state (after about 15 min incubation), where we obtained a high Pearson correlation coefficient (PCC) of 0.78 ± 0.04 (s.d. = standard deviation) and 0.69 ± 0.06 for 0.1 μM and 0.2 μM FtsA respectively **(Fig. 1e)**. We were then wondering how this colocalization would be affected at higher FtsA concentrations. If its localization on the membrane was strictly FtsZ-dependent, we should observe strong colocalization of the two proteins, while excess protein would remain in solution. When we increased the bulk concentration of FtsA to 0.4 and 0.8 μM, we instead found that FtsA pattern abruptly changed to cover the membrane homogeneously, while the FtsZ pattern remained unchanged. Concurrently, the corresponding colocalization coefficient dropped to PCC values of 0.32±0.12 and 0.27±0.09 respectively **(Fig. 1c, 1e)**. This observation suggests that at high concentrations, FtsA binds to the membrane independently of FtsZ filaments. Within this range of FtsA concentrations, we could not observe a significant change in FtsZ treadmilling velocity or monomer residence time **(Fig. S1a, S1b)**. Also the corresponding filament reorganization dynamics as quantified by the decay of the temporal autocorrelation function stayed constant^23^ **(Fig 1i, S1e)**, indicating that in this range of concentrations FtsA does not yet destabilize FtsZ bundles as found previously^21,24^.

Next, we wanted to quantify FtsA-FtsZ co-treadmilling dynamics, i.e. how efficiently FtsZ and FtsA recruit each other to the membrane during filament growth. For this aim, we prepared differential time lapse movies^25^, where we subtract the intensities of consecutive frames to selectively visualize the growing ends of filament bundles (**Fig. 1d)**. We then calculated the Pearson correlation coefficient between the two channels of the differential movies (PCC_diff_), which quantifies the covariation of the fluorescence signals for FtsA and FtsZ at the growing end of a filament bundle with a time resolution of the acquisition rate^4,23,25^. Like the colocalization coefficient (PCC), we found PCC_diff_ to rapidly drop with increasing FtsA:FtsZ ratio, indicating that the ability of FtsZ to dynamically pattern FtsA assemblies on the membrane is severely compromised at high bulk concentrations of FtsA **(Fig. 1d, 1f)**. *In vivo*, this property could contribute to the toxicity of FtsA observed at high expression levels as downstream cell division proteins would bind to FtsA independent of the Z-ring^18,26–28^.

To understand how the activity state of FtsA affects colocalization with FtsZ filaments, we repeated these experiments with FtsA R286W, a well-known ZipA suppressor mutant with decreased self-interaction that is considered to represent an active form of the protein^17^. In contrast to the wild-type protein, we found that this mutant showed more robust colocalization with FtsZ, with high PCC values of around 0.8 at all concentration tested **(Fig. 1e, g)**. We also found FtsA R286W to co-migrate more efficiently with FtsZ filaments with constantly high PCC_diff_ values **(Fig. 1f, h**). At the same time, we could not detect a difference in FtsZ treadmilling velocity, but slightly decreased turnover and increased fluorescence intensity for FtsZ compared to the wildtype protein **(Fig. S1a-d)**. We also found the temporal autocorrelation function decayed more slowly for the filament pattern with FtsA R286W **(Fig. 1i, Fig S1e)**, suggesting that this mutant restricts filament reorganization. Together, these results corroborate earlier *in vivo* observations that FtsA R286W stabilizes the Z-ring^16^ and demonstrate that active FtsA outperforms wildtype FtsA in reproducing the spatiotemporal dynamics of treadmilling FtsZ filaments.

We found that wildtype FtsA and FtsA R286W strongly differ in their ability to localize to the FtsZ filament pattern **(Fig. 1c, g)**. FtsZ interacts with FtsA via a highly conserved C-terminal peptide (CTP)^29^, whose binding site on FtsA is located in its 2B subdomain, close to the Arginine residue mutated in FtsA R286W^9^. Notably, previous yeast two-hybrid experiments suggested that FtsA R286W has an increased affinity towards FtsZ than the wildtype protein^11,17^. To test, if this possibility could explain the better colocalization of FtsA R286W with FtsZ filaments, we measured the binding time of a fluorescently labelled C-terminal FtsZ peptide (TAMRA-KEPDYLDIPAFLRKQAD=TAMRA-CTP) with His-tagged versions of FtsA (FtsA-His and FtsA R286W-His, also see Fig. 4) permanently attached to membranes containing dioctadecylamine (DODA)-tris-NTA, a Ni^2+^-chelating lipid **(Fig. S1f)**. For both versions of FtsA, we found only very transient recruitment of the membrane-bound peptide, with a mean life time of only 90 ± 23ms for FtsA WT and 74 ± 6ms for FtsA R286W (p-value 0.40) **(Fig. 1j, S1g**). A higher affinity of FtsA R286W towards FtsZ monomers seems therefore unlikely to be the reason for its increased colocalization with FtsZ filaments.

Another explanation for the loss of colocalization could be a higher membrane affinity of FtsA WT that results in indiscriminate membrane binding independent of FtsZ. In contrast, an active, monomeric FtsA with low membrane affinity would only be recruited to the membrane in a high avidity complex with FtsZ filaments and predominantly detach from the membrane if not bound to FtsZ. To quantify the intrinsic membrane-affinity of the two versions of FtsA, we used Quartz Crystal Microbalance with Dissipation (QCM-D) and measured the hydrated mass of adsorbed protein on a membrane surface. We found that the membrane affinity of FtsA R286W was only slightly lower than that of FtsA WT (K_d_ 0.32 ± 0.05 μM and of 0.21 ± 0.01 μM respectively) and that the amount of membrane-bound protein saturated at 0.8 μM for both proteins **(Fig. 1k & S1h)**. This result is consistent with the observation that FtsA WT and FtsA R286W behave identical in co-sedimentation experiments^10^. This small difference in membrane binding affinities to the membrane cannot explain the observed contrast in colocalization of the two version of FtsA with FtsZ filaments in particular at high bulk concentrations of the proteins.

Wildtype FtsA has the tendency to form membrane-bound arrays of minirings, while FtsA R286W was found to assemble into tightly packed short filaments and arcs^10^. Accordingly, enhanced colocalization with FtsZ could be because of a higher packing density of FtsA R286W below FtsZ filaments. To test this hypothesis, we analysed the pixel-by-pixel relationship between the fluorescence intensities of FtsZ and FtsA in dual-colour fluorescence time lapse movies. The linear slope of this relationship shows how the density of FtsA on the membrane changes when the FtsZ filament density increases and therefore is an indicator for the binding capacity of FtsZ filaments for FtsA **(Fig. S1i)**. At low FtsA concentrations, we found the slopes for both proteins to be similar. Above 0.2 μM these values dropped significantly for FtsA WT, but remained constant for FtsA R286W at ∼0.7 **(Fig. 1l, S1i)**. These results indicate that at high concentrations the amount of FtsA WT that can be recruited to FtsZ filaments is strictly limited, likely due to the formation of minirings, while FtsA R286W continues to accumulate.

### FtsA R286W allows for enhanced recruitment of FtsN_cyto_ to FtsZ filaments

FtsA provides a physical link between treadmilling FtsZ filaments in the cytoplasm and cell division proteins located in the membrane. However, polymerization of FtsA and recruitment of downstream proteins are thought to be mutually exclusive as they both involve interactions via FtsA’s 1C domain^30^. Accordingly, in an active, depolymerized FtsA this domain would be readily available to interact with downstream proteins like FtsN. Vice versa, binding of these proteins should facilitate the transition of FtsA from the inactive, oligomeric to the active, more monomeric form.

To test these predictions, we mimicked the presence of transmembrane FtsN in the bilayer by attaching its His-tagged, cytoplasmic peptide (FtsN^1-32^-His6x=FtsN_cyto_) to the surface of a supported membrane containing 0.25% Tris-NTA lipids^4^. We then used these modified membranes to compare how the two versions of FtsA differ in their ability to recruit FtsN_cyto_ to treadmilling filaments of FtsZ **(Fig. 2a)**. Closely mirroring the behaviour observed for the localizations of FtsA (**Fig. S2a, b)**, we found strong overlap (PCC) and co-treadmilling (PCC_diff_) of FtsN_cyto_ with FtsZ filaments at low concentrations of FtsA WT ([FtsA WT] < 0.4 μM) and a sudden drop of these values at higher concentrations **(Fig. 2b-d)**. In contrast, in the case of FtsA R286W, both values remained higher than 0.6, even at concentrations above 0.4 μM **(Fig. 2c-e)**.

**Figure 2:**
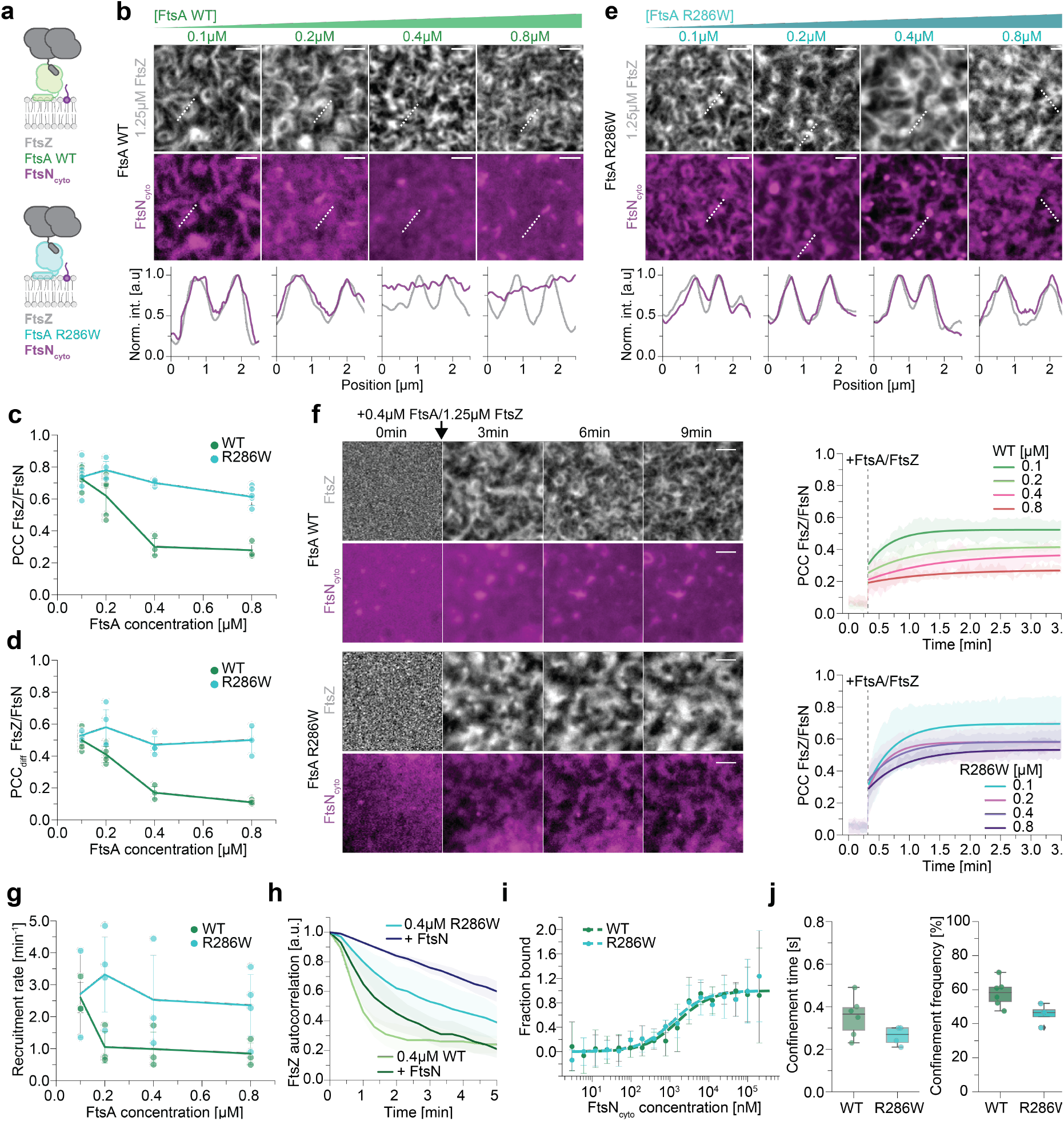
FtsA R286W shows enhanced recruitment of FtsN_cyto_ to FtsZ filaments. **a**, Cartoon illustrating which components are present in the TIRF experiment (**b** and **e)** and which are labeled (= bold text). **b**, Representative micrographs of A488-FtsZ (grey) and Cy5-FtsN_cyto_ (magenta) at increasing FtsA WT and constant FtsZ concentration. FtsN_cyto_ colocalizes well with FtsZ filaments up to 0.2 μM FtsA WT, but fails at higher concentrations. The line profiles correspond to the dashed white lines. **c**, Colocalization of FtsZ/FtsN quantified by PCC, shows that colocalization of FtsZ with FtsN differs significantly at FtsA concentrations above 0.2 μM (FtsA WT: 0.30 ± 0.05; FtsA R286W: 0.70 ± 0.02; p-value: 4.92*10^−4^). **d** Dynamic colocalization of FtsZ/FtsN quantified by ΔPCC. FtsN_cyto_ follows FtsZ treadmilling more efficient with FtsA R286W at concentrations above 0.2 μM (0.17 ± 0.04; 0.47 ± 0.05; p-value: 3.79*10^−3^). **e**, Representative micrographs of A488-FtsZ (grey) and Cy5-FtsN_cyto_ (magenta) at increasing FtsA R286W and constant FtsZ concentration. FtsN_cyto_ colocalizes well with FtsZ at all tested FtsA R286W concentrations. The line profiles correspond to the dashed white lines. **f**, Top: From a homogeneous distribution, Cy5-FtsN_cyto_ (magenta) does not colocalize with FtsZ filaments on the membrane after the addition of 1.25 μM A488-FtsZ (grey) and 0.4 μM FtsA WT at 0 min. Bottom: The same experiment with 0.4 μM FtsA R286W reveals that Cy5-FtsN_cyto_ colocalizes well with FtsZ filaments after protein addition at 0 min. Right: Mean values of PCC versus time at different FtsA WT and R286W concentrations. FtsN_cyto_ is recruited to FtsZ filaments at all tested FtsA R286W concentrations, but the PCC remains low at FtsA WT concentrations above 0.2 μM. **g**, Quantification of the recruitment rate of FtsN_cyto_ towards FtsZ filaments, extracted from experiments in **f** by fitting a power law exponential. FtsA R286W (cyan) recruits FtsN_cyto_ consistently fast at all concentrations, whereas the recruitment rate decreases for FtsA WT (green) above 0.2 μM (1.05 ± 0.49; 3.32 ± 1.17; p-value: 0.045). **h**, Presence of FtsN_cyto_ increases the persistency of the FtsZ network drastically. This effect is observed with both FtsA variants. **i**, Quantification of the binding affinity of FtsN_cyto_ towards FtsA WT or R286W by MST shows that there is no difference (1.58 ± 0.43 mM and 1.17 ± 0.37 mM respectively). **j**, Quantification of FtsN_cyto_ confinement events to FtsA/FtsZ cofilaments by single molecule tracking. While the confinement period for FtsN_cyto_ is slightly, but not significantly longer in the presence of FtsA WT (0.35 ± 0.09s vs. 0.26 ± 0.04s) the confinement frequency is increased significantly for FtsA WT (57.99 ± 7.29%; 45.70 ± 5.10; p-value: 0.03).

Next, we were wondering about the rate of FtsN_cyto_ accumulation on FtsZ-FtsA co-filaments. Starting from a homogeneous distribution of the membrane-bound peptide, we measured how quickly the overlap of the FtsN_cyto_ and FtsZ signals increased after adding FtsA and FtsZ **(Fig. 2f, S2c,d)**. By fitting an exponential function to the increase of the PCC with time, we were able to extract the corresponding recruitment rate **(Fig. 2f,g)**. While FtsN enrichment saturated within 1 min for all concentrations of FtsA R286W, the recruitment rate for FtsA wt dropped significantly already at 0.2 μM FtsA such that it required more than twice as long for FtsN to colocalize with FtsZ. Together, these data demonstrate FtsA R286W recruits downstream proteins to FtsZ filaments more efficiently than wildtype FtsA. This property could also explain why cell division is faster in cells with FtsA R286W^31^.

Previous literature suggested that arrival of FtsN at the Z-ring triggers disassembly of FtsA WT oligomers into a more FtsA R286W-like, monomeric state^7^. Thus, we were wondering if the presence of FtsN_cyto_ could change the colocalization of FtsA WT with FtsZ to resemble the behaviour of FtsA R286W. While addition of FtsN_cyto_ slightly increases the total amount of both proteins on the membrane, their overlap and the density of FtsA on FtsZ filaments, it could not prevent the loss of colocalization with FtsZ at higher concentrations of FtsA **(Fig. S2e-h)**, indicating that the presence of FtsN_cyto_ has no strong effect on the recruitment of FtsA towards FtsZ filaments. Interestingly, we found that adding FtsN_cyto_ significantly slows down the reorganization dynamics of the FtsZ pattern, suggesting that binding of FtsN_cyto_ leads to a transition in FtsA that prevents FtsZ filament realignment **(Fig. 2h, S2i)**. However, FtsN_cyto_ alone is not sufficient to fully convert FtsA WT into FtsA R286W.

Next, we wanted to know if the increased overlap between FtsN_cyto_ and FtsZ with FtsA R286W can be explained by an increased affinity of FtsN towards the hypermorphic mutant as suggested previously^5,17^. To test this idea, we performed microscale thermophoresis experiments (MST) with fluorescently labelled FtsA WT and FtsA R286W and increasing concentrations of FtsN_cyto_ **(Fig. 2i and S2j)**. For both proteins, we measured similar dissociation constants in these experiments of *K*_*D*_(wt Cy5-FtsA/FtsN_cyto_) = 1.58 ± 0.43 μM and *K*_*D*_(Cy5-FtsA R286W/FtsN_cyto_) = 1.17 ± 0.37 μM. Since FtsN interacts with FtsA on the membrane surface, we were wondering if two-dimensional confinement could enhance a difference. We therefore imaged the trajectories of individual membrane-bound FtsN_cyto_ peptides in the presence of treadmilling FtsZ-FtsA filaments and then quantified the duration and frequency of confinement^4^ **(Fig. 2j, S2k, l)**. We found that both values were in fact slightly lower for FtsA R286W confirming that it does not have increased affinity towards FtsN_cyto_.

We conclude that our *in vitro* experiments recapitulate several observations made in the living cell, where FtsA R286W recruit FtsN to FtsZ filaments more efficiently and its arrival at midcell stabilizes the Z-ring. However, we found this difference is not due to an enhanced affinity of this peptide to FtsA R286W, but likely the result of the higher packing density of FtsA R286W below FtsZ filaments **(Fig. 1l**).

### FtsA R286W shows faster membrane exchange than FtsA WT, while the self-interaction of both proteins is enhanced in the presence of FtsN_cyto_

So far, we have seen that FtsA R286W is more strongly recruited to FtsZ filaments **(Fig. 1**), which allows for an improved recruitment of FtsN_cyto_ towards FtsA-FtsZ co-filaments **(Fig. 2**). Additionally, we also found that FtsA R286W co-migrates with treadmilling FtsZ filaments more efficiently **(Fig. 1d, f, h)**. As FtsA WT and FtsA R286W do not significantly differ in their affinities towards FtsZ **(Fig. 1j)**, the membrane **(Fig. 1k)** or FtsN_cyto_ **(Fig. 2i, j)**, we decided to investigate a possible mechanism that could explain our observations.

First, we studied the behaviour of single FtsA proteins on the membrane surface. In a background of unlabelled proteins, we followed individual FtsA WT and FtsA R286W proteins and analyzed their trajectories by single particle tracking **(Fig. 3a)**. At 0.1μM, we found that FtsA WT showed a low mobility with a diffusion constant of 0.14 ± 0.04μm^2^/s and a mean residence time of 10.2 ± 0.7 s. With increasing bulk concentrations and protein densities on the membrane, the mean residence time remained constant (9.39 ± 0.28s at 0.8 μM FtsA), while the diffusion constant dropped to a value of 0.004 ± 0.001μm^2^/s indicating almost immobile proteins on the membrane. Similar to previous FRAP experiments *in vivo*^16^, we found a faster exchange of FtsA R286W with a single molecule residence time 2-10 fold shorter than that of wildtype FtsA (from 0.54 ± 0.14s to 4.8 ± 0.48s for 0.1 and 0.8 μM respectively). The diffusivity of FtsA R286W also decreased at higher protein concentrations (from 0.42 ± 0.06 μm^2^/s to 0.054 ± 0.011 μm^2^/s for 0.1 and 0.8 μM respectively), but remained mobile even at higher concentrations **(Fig. 3b)**. These differences in residence time and diffusion likely correlate with different modes of self-interaction of the two proteins^10,17^.

**Figure 3:**
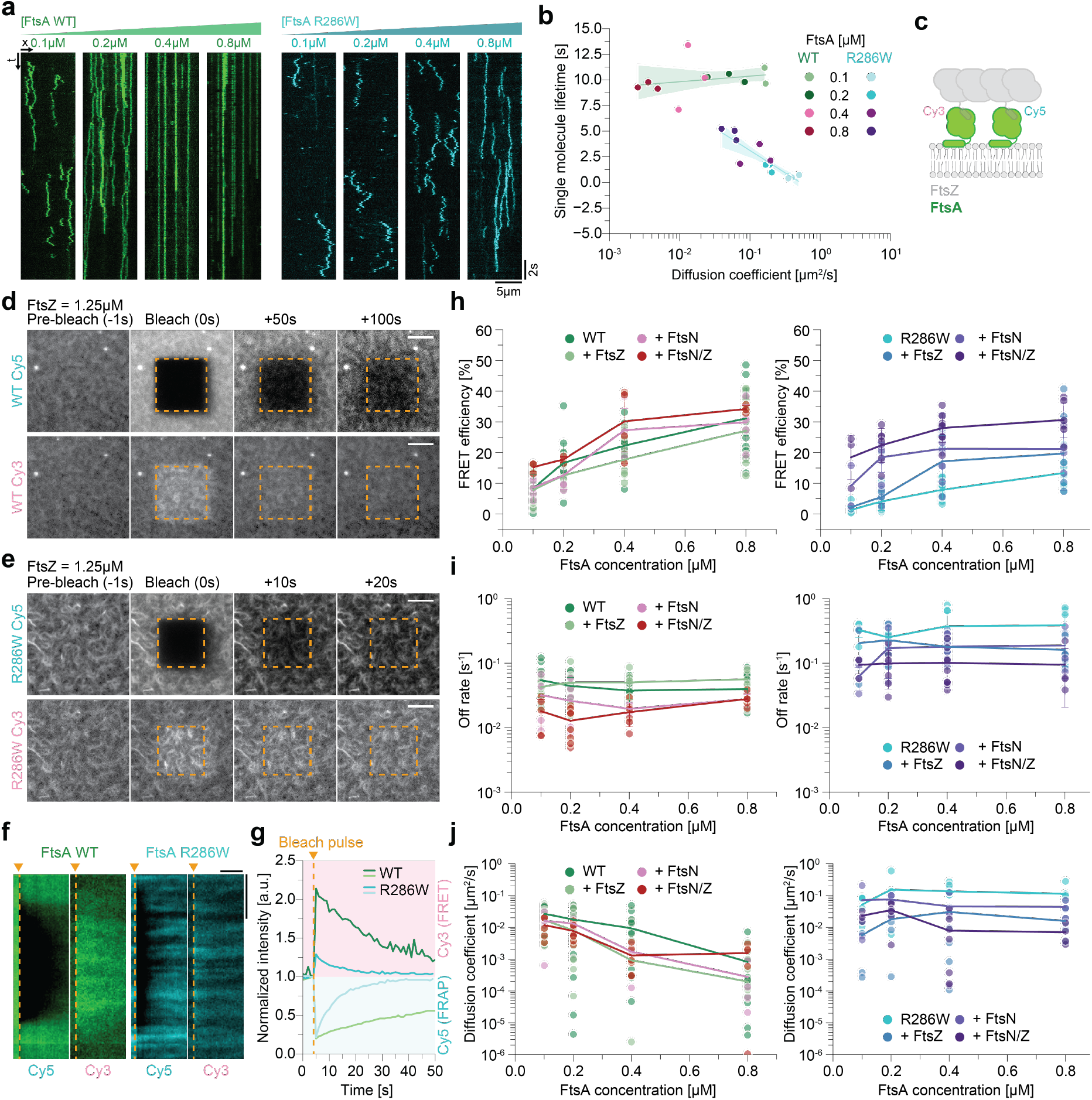
FtsA R286W shows faster membrane exchange than FtsA WT, while the self-interaction of both proteins is enhanced in the presence of FtsN_cyto_. **a**, Representative kymographs of single molecules of Cy5-FtsA WT (green) and Cy5-FtsA R286W (cyan) at increasing concentrations. FtsA WT remains immobile above 0.2 μM, while FtsA R286W slows down, but displays diffusive behavior at all concentrations shown. **b**, The diffusion coefficient of FtsA WT decreases with increasing concentrations (0.14 ± 0.04 μm^2^/s to 0.004 ± 0.0009 μm^2^/s) while its lifetime remains stable (10.21 ± 0.70 s to 9.40 ± 0.28 s). The diffusion coefficient of FtsA R286W also decreases, but is still 10x higher than for FtsA WT (0.41 ± 0.06 μm^2^/s to 0.05 ± 0.01 μm^2^/s). The lifetime also increases, but remains lower compared to FtsA WT (0.54 ± 0.14 s vs. 4.80 ± 0.49 s). **c**, Schematic of the experiment to measure FRET between FtsA labeled either with Cy3 or Cy5. **d**, Representative micrographs of FRET assay performed by acceptor photobleaching (Cy5-FtsA WT) in the presence of the donor Cy3-FtsA WT, mixed in the ratio 1:1. Top: The fluorescence signal of Cy5-FtsA WT recovers within 100s. Bottom: The corresponding increase in Cy3-FtsA WT intensity is strong and long-lived. **e**, Representative micrograph of an acceptor photobleaching experiment of Cy3/Cy5-FtsA R286W + FtsZ. Top: The fluorescence signal of Cy5-FtsA R286W recovers within 20s. Bottom: The corresponding increase in Cy3-FtsA R286W intensity is weak. Scale bars in **d** and **e** are 5 μm. **f**, Kymographs depicting the differences in FRAP recovery (left), as well as the duration of the FRET signal (right) for FtsA WT (green) and FtsA R286W (cyan). Scale bars are 4 μm and 20 sec, respectively. **g**, Representative examples of FRAP recovery (bottom, grey rectangle) and FRET increase curves (top, pink rectangle) for FtsA WT (green) and R286W (cyan). **h**, Left: FRET of FtsA WT increases at higher concentrations (from 8.11 ± 4.43 % to 30.63 ± 9.86 %). Addition of FtsZ slightly decreases measured FRET efficiency, but not significantly (22.14 ± 5.73% for WT alone vs. 17.74 ± 7.31% in FtsZ presence, p-value: 0.17 at 0.4μM). The presence of FtsN_cyto_ and FtsZ/FtsN_cyto_ increases the self-interaction (22.14 ± 5.73% vs 27.29 ± 7.13% with FtsN_cyto_ and 30.17 ± 9.64% with FtsZ/FtsN_cyto_ at 0.4 μM, p-values: 0.16 and 0.05). Right: FtsA R286W FRET also increases with concentration, but is constantly significantly lower than for FtsA WT (1.35 ± 1.19 % to 13.36 ± 4.59 %). FtsZ increases FRET slightly, while FtsN_cyto_ drastically increases measured FRET (8.09 ± 2.79% vs 18.19 ± 8.78% with FtsZ and 21.26 ± 6.98% with FtsN_cyto_ at 0.4μM; p-values: 1.4*10^−3^ and 2.03*10^−5^ respectively). With both, FtsZ and FtsN_cyto_, the FRET signal of WT and R286W are indistinguishable (30.17 ± 0.9.64% vs. 28.01 ± 4.62% at 0.4 μM). **i**, Left: Off-rates remain constantly slow with increasing concentrations of FtsA WT (0.054 ± 0.025 s^-1^ to 0.039 ± 0.019 s^-1^). Presence of FtsN and FtsZ/FtsN decreases the off-binding rate of FtsA WT (0.038 ± 0.012s^-1^ vs. 0.019± 0.004s^-1^ and 0.017± 0.007s^-1^ at 0.4 μM; p-values: 6.5*10^−3^ and 1.55*10^−3^). Right: Off-binding rates remain constantly fast with increasing concentrations of FtsA R286W (0.33 ± 0.08 s^-1^ to 0.39 ± 0.26 s^-1^). FtsN and FtsZ/FtsN decrease off-binding rate of FtsA R286W (0.37 ± 0.24s^-1^ vs. 0.179± 0.07s^-1^ and 0.10± 0.06s^-1^ at μM FtsA R286W; p-values: 0.1 and 0.007). However, FtsA R286W is still more dynamic than FtsA WT (0.02 ± 0.006 s^-1^ vs. 0.10 ± 0.074 s^-1^ at 0.8 μM with FtsZ/N). **j**, Left: Diffusion coefficient drops with increasing concentrations of FtsA WT (0.027 ± 0.018 μm^2^/s to 0.0008 ± 0.0017 μm^2^/s). Additional components slightly decrease the mobility (0.009 ± 0.01μm^2^/s vs 0.0009 ± 0. 0008 μm^2^/s and 0.0013 ± 0.001 μm^2^/s for FtsA WT and with FtsN_cyto_ or FtsZ/FtsN_cyto_ respectively, p-values: 0.22 and 0.16). Right: Diffusion coefficient remains unchanged at increased concentrations of FtsA R286W (0.052 ± 0.034 μm^2^/s to 0.114 ± 0.07 μm^2^/s). The presence of FtsN, FtsZ individually and together slow down diffusion of FtsA R286W (0.14 ± 0.07μm^2^/s vs 0.046 ± 0. 036 μm^2^/s and 0.0079 ± 0.01 μm^2^/s for FtsA R286W alone and with FtsN_cyto_ or FtsZ/FtsN_cyto_ respectively, p-values: 0.03 and 3.25*10^−4^). However, D_coef_ remains still higher than for FtsA WT (0.0015 ± 0.0011 μm^2^/s vs. 0.007 ± 0.0043 μm^2^/s at 0.8 μM with FtsZ/N).

To directly measure FtsA self-interaction in our fluorescence microscopy experiments, we established a FRET (Foerster resonance energy transfer)-based assay using FtsA labelled with either Cy5 or Cy3. First, we tried to find evidence for self-interaction in solution, but could not detect any significant FRET signal under these conditions or oligomerization in SEC-MALS experiments **(Fig. S3a, S3b)** demonstrating that FtsA is monomeric in solution. However, we found that FRET increased significantly in the presence of lipid vesicles, indicating that oligomerization of FtsA depends on the interaction with lipid membrane **(Fig. S3b-S3c)**. We then decided to use this approach to measure the degree of FtsA self-interaction on supported lipid membranes by quantifying the change in donor fluorescence after photobleaching the acceptor fluorophore^32–34^ **(Fig. 3c-e and Fig. S3d)**. As a negative control, we attached His-SUMO labeled with these fluorophores to membranes with an increasing fraction of Tris-NTA lipids. For this negative control, we did not find any significant FRET, even at maximal coverage of the membrane surface **(Fig. 4g)**. We also saw no change in donor intensity in photobleaching experiments without acceptor fluorophore. **(Fig. S3e)**.

**Figure 4:**
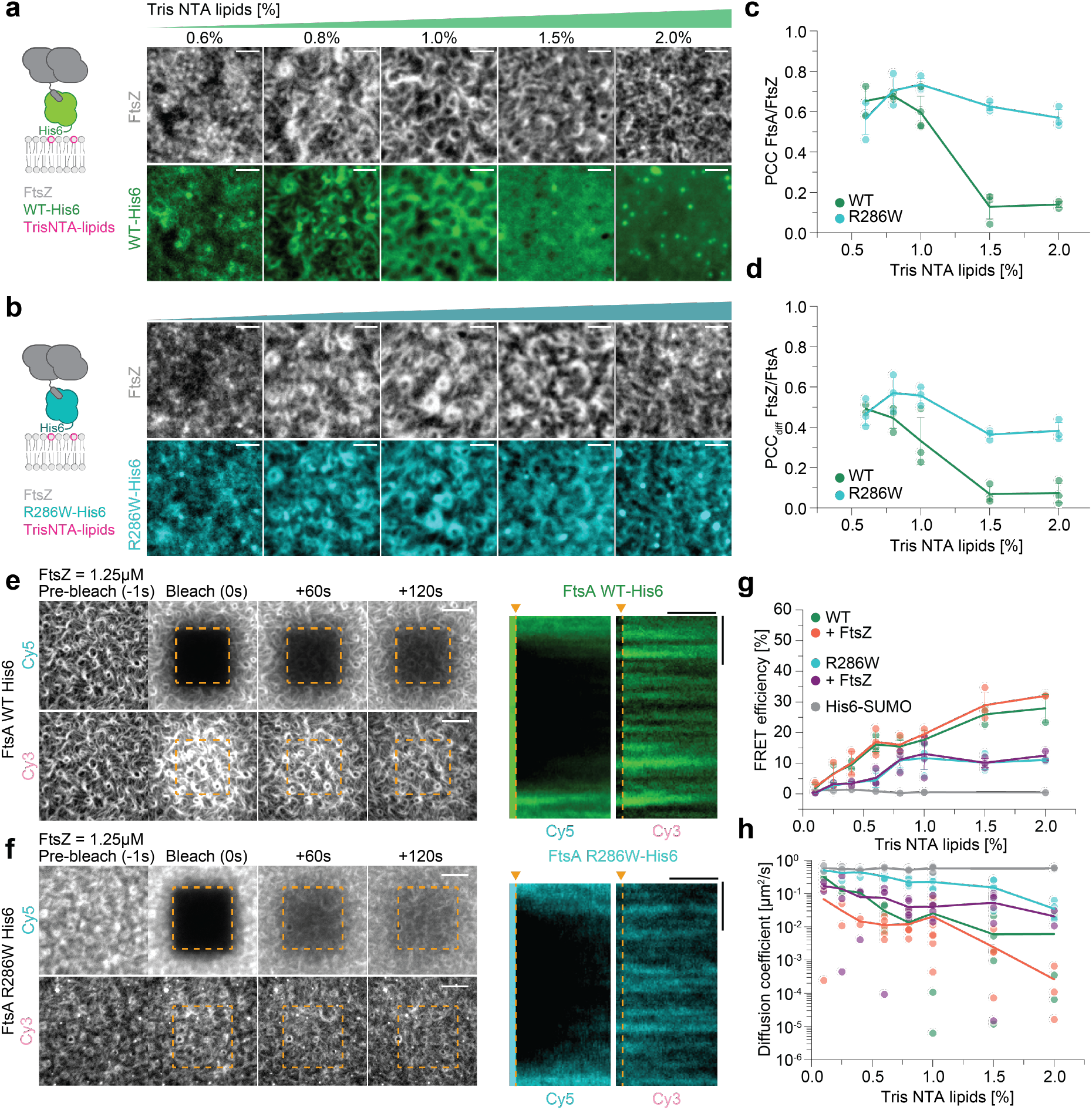
Membrane diffusion and protein binding dynamics contribute to co-treadmilling of FtsA with treadmilling filaments of FtsZ. **a**, Representative micrographs of A488-FtsZ (grey) and Cy5-FtsA WT-His6 (green) at increasing Tris-NTA lipid concentrations. With more than 1% Tris-NTA lipids FtsA WT-His6 does not colocalize to FtsZ filaments. **b**, Representative micrographs of A488-FtsZ (grey) and Cy5-FtsA R286W-His6-(cyan) at increasing Tris-NTA concentrations. **c**, Colocalization of FtsA-His6/FtsZ quantified by PCC. Above 1% Tris-NTA lipids the colocalization of His-tagged FtsAs with FtsZ differs significantly (0.13 ± 0.06; 0.63 ± 0.02; p-value: 3.86 * 10^−4^). **d**, Dynamic colocalization FtsA-His6/FtsZ quantified by PCC_diff_. With more than 1% Tris-NTA lipids the co-treadmilling of His-tagged FtsAs with FtsZ differs significantly (0.07 ± 0.04; 0.36 ± 0.02; p-value: 5.70 * 10^−4^). **e**,**f**, Representative micrographs of acceptor bleaching recovery and donor intensity increase of FtsA WT-His6 (**e**) and FtsA R286W-His6 (**f**). Right: Kymographs depicting the FRAP recovery, as well as the duration of the FRET signal for FtsA WT-His6 (green) and FtsA R286W-His6 (cyan). **g**, FRET of His-tagged FtsAs and the His6-SUMO control. FRET for His-SUMO is not detected, whereas His-tagged FtsAs self-interaction rises with increasing protein density. FRET signal of FtsA WT is significantly higher compared to FtsA R286W-His6. The presence of FtsZ has no significant effect on the measured self-interaction. **h**, Diffusion coefficient of His-tagged FtsAs ± FtsZ and the His6-SUMO control. While the mobility of His6-SUMO remains constant, the diffusion of His-tagged FtsAs decreases at higher protein density.

We found that the FRET efficiency for FtsA WT was generally higher than for R286W **(Fig. 3c-3h)**, and that it increased with the bulk concentration indicative for stronger FtsA WT self-interaction. We could also use the data from these bleaching experiments to quantify the membrane binding kinetics and diffusion of membrane-bound proteins by analysing the change of the fluorescence profile during recovery^35^. In agreement with our single molecule experiments, we found a much faster exchange **(Fig. 3i)** and diffusion for FtsA R286W compared to the wildtype protein **(Fig. 3j)**. This fast turnover of FtsA R286W on the membrane can also explain the shorter confinement time we found for FtsN_cyto_ (**Fig. 2j**). Consistent with our QCM-D experiments **(Fig. 1k)**, we found that FRET and membrane exchange plateaued at FtsA concentrations above 0.8 μM **(Fig. S3f-S3h)**. Furthermore, we found that replacing ATP with a non-hydrolysable analogue ATPγS had no significant effect on membrane-binding, self-interaction or membrane binding dynamics **(Fig. S3i-S3k)**. Together, these data confirm that self-interaction of membrane-bound FtsA WT is enhanced, which results in slower exchange dynamics compared to the mutant protein.

Next, we wanted to find out how binding of FtsZ and FtsN_cyto_ affect FtsA protein exchange, diffusion and self-interaction on the membrane. In the presence of FtsZ, FtsA 286W is still exchanged one order of magnitude faster than the wildtype protein, with membrane off-rates of around 0.20 s^-1^ and 0.05 s^-1^ respectively **(Fig. 3i)**. As the off-rates for FtsZ are between 0.10 and 0.20 s^-1^ **(Fig. S1b,c)**, this means that FtsA R286W turns over 1-2 times faster than FtsZ monomers in the treadmilling filament, while FtsA WT remains bound about 2-4 times longer. For both versions of FtsA, the diffusion coefficient was decreased in the presence of FtsZ **(Fig. 3j)**. Next, we were interested in testing if FtsN_cyto_ promotes de-oligomerization of FtsA WT and reduces the corresponding FRET efficiency^7^. Surprisingly, we found that for FtsA WT the presence of FtsN_cyto_ resulted in a small FRET increase. In the case of FtsA R286W we found a reduced, albeit present FRET signal, which was further increased three-fold when both binding partners, FtsN_cyto_ and FtsZ were present. In addition, we found that the presence of FtsN_cyto_ slightly decreased the detachment rate and membrane mobility of both FtsA WT and FtsA R286W **(Fig. 3i-j)**. Together, these observations show that in contrast to previous reports, FtsN_cyto_ does not trigger disassembly of FtsA oligomers^3,14,17^. Instead, our data suggests that FtsN_cyto_ supports the formation of a distinct, oligomeric structure of FtsA, possibly to due enhanced lateral interactions^10^, which results in a higher FRET efficiency for both versions of FtsA.

### Lateral diffusion and membrane binding dynamics contribute to co-treadmilling of FtsA with treadmilling FtsZ filaments

Co-treadmilling of FtsA with FtsZ relies on the dynamic exchange of FtsA on the membrane. We have found that FtsA R286W shows a dramatically shorter residence time and decreased self-interaction compared to the wildtype protein **(Fig. 3)**. These two properties strongly correlate with a much stronger colocalization with treadmilling FtsZ filaments **(Fig. 1)**. To investigate the respective contributions of membrane binding kinetics and FtsA self-interaction on the colocalization dynamics with FtsZ, we wanted to create variants of the two FtsA proteins that are permanently attached to the membrane. We therefore replaced their amphipathic helices by C-terminal His-tags to obtain FtsA WT(1-405)-6xHis (=FtsA-His) and FtsA R286W (1-405)-6xHis (=FtsA R286W-His), which we attached to membranes containing different amounts of Tris-NTA lipids. Accordingly, in these experiments the proteins only differ in their tendency to form oligomers.

When we measured the colocalization of FtsZ with the His-tagged versions of FtsA, we found that just like for the native proteins, colocalization and co-treadmilling decreased with increasing FtsA-His densities on the membrane, while they stayed almost constant for FtsA R286W-His **(Fig. 4a-d)**. Furthermore, despite identical densities on the membrane at a given amount of Tris-NTA lipids, colocalization of FtsA R286W-His with FtsZ was always higher than for FtsA WT-His, confirming that FtsZ filaments have a higher capacity for FtsA R286W than for the wildtype protein^10^ **(Fig. 1l)**. Interestingly, co-treadmilling with FtsZ was about 50% reduced for FtsA R286W-His compared to the reversibly membrane-binding protein **(Fig. 1f)**, emphasizing that although the fast diffusion of FtsA R286W-His allows for some degree of co-treadmilling with FtsZ filaments, recruitment of the protein from solution significantly contributes to its efficiency.

To evaluate the degree of FtsA self-interaction in the absence of protein exchange, we measured the FRET efficiency and diffusion constant of FtsA-His and FtsA R286W-His at different densities. Similar to the proteins with native membrane binding, His-tagged FtsA R286W showed lower FRET efficiency and faster diffusion at all densities tested **(Fig. 4e-h and Fig. S4c-S4e)**. In addition, as the FRET efficiency for permanently attached FtsA R286W-His was not higher than for the reversibly binding protein, we can conclude that faster membrane-binding dynamics are not a limiting factor for FtsA self-interaction. Conversely, the different FRET efficiencies and membrane mobilities we observe for FtsA-His and FtsA R286W-His are solely due to the proteins existing in different oligomeric states.

### A model for the behaviour of FtsA during divisome maturation

Using a minimal set of purified components in combination with quantitative fluorescence microscopy, we were able to provide new insights into the role of FtsA during the assembly and activation of the bacterial cell division machinery. We confirmed previous conclusions on the properties of FtsA and its hyperactive mutant FtsA R286W based on *in vivo* observations. First, we demonstrate that FtsZ filaments define the spatiotemporal distribution of FtsA WT assemblies on the membrane^19^ and that FtsA R286W stabilizes FtsZ filament reorganization compared to the wildtype protein^8^ **(Fig. 1)**. Second, we confirm that FtsA R286W outperforms FtsA WT in the recruitment of downstream proteins^17^ **(Fig. 2)**. Furthermore, our results from FRET and single-molecule experiments support previous findings on the oligomeric nature of FtsA as well as a decreased self-interaction in case of FtsA R286W^17^ **(Fig. 3)**.

Importantly, our experiments revealed that binding of FtsN_cyto_ does not disassemble FtsA WT oligomers. Instead, we find that the presence of FtsN_cyto_ increases FtsA self-interaction in particular for FtsA R286W. Previous electron microscopy studies found that FtsA is able to form oligomers of different conformations, such as straight filaments and minirings, but also filament doublets, which form predominantly in the case of several ZipA suppressor mutants^9,10,36^. It therefore seems likely that FtsA can oligomerize via different interfaces allowing for lateral and longitudinal interactions^10^. As longitudinal interactions are compromised in the case of FtsA R286W^10,36^, binding of FtsN_cyto_ likely induces a conformational change that promotes lateral interactions, enhancing the formation of filament doublets and increasing the FRET efficiency **(Fig. 3h, 5c)**. This FtsN-dependent structural transition of FtsA goes along with an enhanced recruitment of FtsZ filaments and a decrease of filament reorientation **(Fig. 2h, S2f)**. Importantly, these interpretations are supported by a concurrent study that finds that FtsA switches from minirings to double filaments upon binding of FtsN (Jan Löwe, MRC LMB, Cambridge, personal communication).

The concentration dependent tendency of FtsA to organize into ring arrays^10^ also offers an explanation for the transition from co-migrating dynamic assemblies towards a stable, homogeneous protein layer on the membrane **(Fig. 1c, Fig. 3a, Fig. 5a**,**b)**. Conversely, the absence of rings in case of FtsA R286W can account for its faster membrane exchange as well as the higher packing density below FtsZ filaments **(Fig. 1g, 3a, 5c)**. Importantly, the increased density of FtsA R286W leads to an enhanced recruitment of FtsN_cyto_ **(Fig. 2)** and possibly other downstream proteins with weak affinities towards FtsA such as FtsQ^4,6,37^ and FtsW^12^. This property could contribute to the ability of FtsA R286W to bypass otherwise essential cell division proteins^17^.

**Figure 5:**
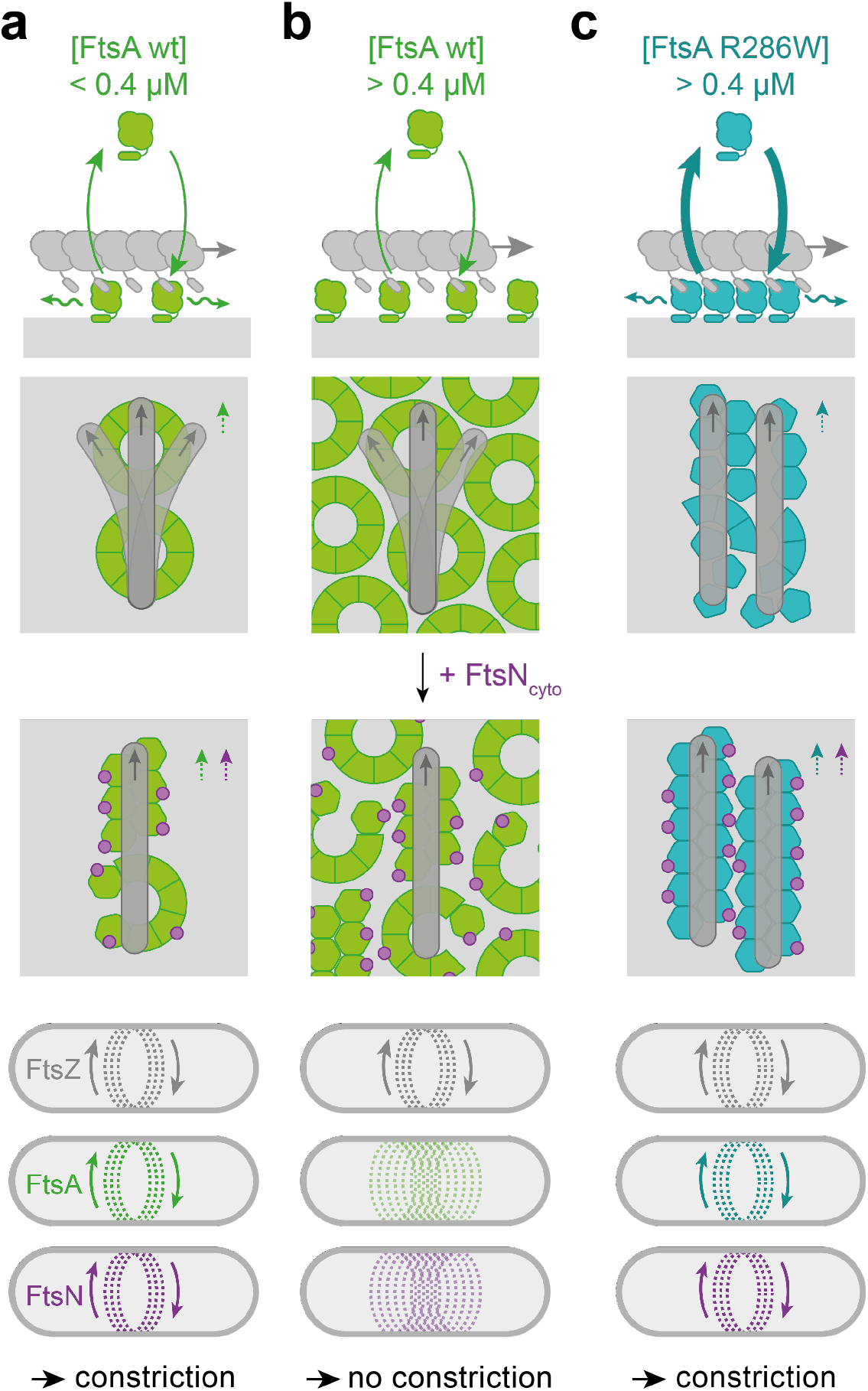
A model for the behavior of FtsA during divisome activation. FtsA and FtsZ form co-treadmilling filaments on the membrane surface. The spatiotemporal distribution of FtsA on the membrane relies on its dynamic interaction with FtsZ filaments and the lipid membrane as well as its exchange by lateral diffusion. Grey arrows indicated treadmilling direction, curved arrows the dynamic exchange of FtsA to and from the membrane, wavy arrows represent lateral diffusion, dashed arrows indicate co-treadmilling of FtsA and FtsN. **a**, At low FtsA WT concentrations, i.e. at a protein ratio found *in vivo* ([FtsA WT] = 0.2 μM, [FtsZ] = 1.25 μM), fast diffusion of membrane-bound FtsA allows for co-treadmilling with FtsZ filaments despite slow cycling on and off the membrane. The presence of FtsA minirings limits the packing density and therefore the amount of FtsA recruited to the filament. Furthermore, it allows for a continuous realignment of treadmilling filaments. Recruitment of FtsN_cyto_ does not depolymerize FtsA oligomers, but supports a conformational change that prevents reorganization of filaments. While FtsZ, FtsA and downstream division proteins are dynamically moving around the cell, they distribute cell wall synthesis allowing for cell constriction. **b**, At higher concentrations, FtsA forms a continuous array of minirings on the membrane, stabilized by lateral interactions. Due to the absence of diffusion, FtsA cannot follow FtsZ treadmilling dynamics and fails to distribute cell wall synthesis. As a result, the cell fails to divide. **c**, Even at high concentrations of FtsA R286W, its fast exchange allows for a close replication of FtsZ filament dynamics on the membrane surface. Loss of longitudinal interactions in this mutant disrupts minirings, allowing for a higher packing density of FtsA which limits the realignment of filaments. Binding of FtsN enhances lateral interactions to promote a structural change that further limits realignment of filaments. As all proteins are dynamically moving around the cell, constriction and cell division is possible even at elevated levels of FtsA R286W.

Finally, our experiments also shed light on the signalling function of FtsA during cytokinesis, i.e. its ability to transmit the spatiotemporal information originating from FtsZ filaments in the cytoplasm towards the periplasmic space. This function relies on a close replication of FtsZ polymerization dynamics by FtsA on the membrane surface, which is the result of its dynamic exchange by lateral diffusion and via membrane binding and detachment. Both of these two processes are much faster for FtsA R286W than for FtsA WT **(Fig. 3i, j**). As FtsA R286W turns over up to two times faster than FtsZ, this membrane anchor can sample the dynamic filament with a minimal loss of spatiotemporal information^38^. Together, we expect FtsA R286W to not only be better at directing cell division to midcell, but also at homogenously distributing cell wall synthesis around the division site. It will be interesting to study the behavior of single FtsA molecules *in vivo* and how different exchange dynamics of the membrane anchor correlate with the ability of treadmilling FtsZ filaments to drive the directional motion of cell division proteins^39^.

With the described benefits of the R286W mutation for the roles of FtsA, the question arises why it did not persist during evolution. *In vivo*, FtsA R286W produces misaligned and twisted division septa and minicells. Accordingly, it is possible that the longer residence times of FtsA WT oligomers, a tighter control of their hypothesized structural reorganization, as well as the dependency on ZipA as a second membrane anchor provide additional control mechanisms that increase the precision and robustness of cell division.

## Acknowledgments

We acknowledge members of the Loose laboratory at IST Austria for helpful discussions—in particular L. Lindorfer for his assistance with cloning and purifications. We thank J. Löwe and T. Nierhaus (MRC-LMB Cambridge, UK) for sharing unpublished work and helpful discussions, as well as D. Vavylonis and D. Rutkowski (Lehigh University, Bethlehem, PA, USA) as well as S. Martin (University of Lausanne, Switzerland)) for sharing their code for FRAP analysis. We are also thankful for the support by the Scientific Service Units (SSU) of IST Austria through resources provided by the Bioimaging Facility (BIF) and the Lab Support Facility (LSF). This work was supported by the European Research Council through grant ERC 2015-StG-679239 and by the Austrian Science Fund (FWF) StandAlone P34607 to M.L. and HFSP LT 000824/2016-L4 to N.B. For the purpose of open access, we have applied a CC BY public copyright licence to any Author Accepted Manuscript version arising from this submission.’

## Supplementary Figures

**Figure S1:**
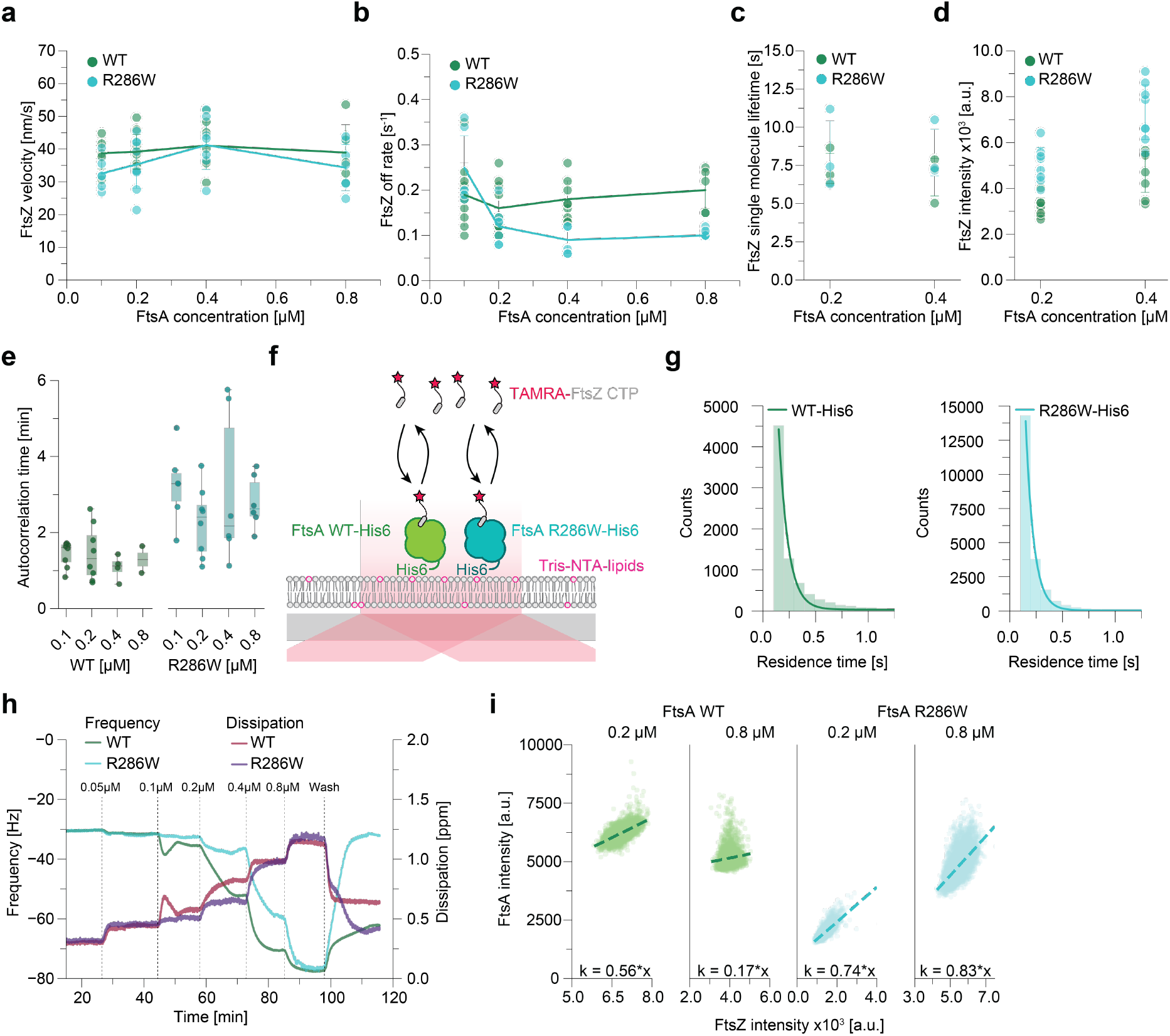
FtsZ dynamics are identical with FtsA WT and R286W, more FtsZ is recruited by FtsA R286W. **a**, FtsZ treadmilling velocity is identical in the presence of FtsA WT or R286W at all concentrations tested (38.95 ± 8.51nm/s vs. 34.35 ± 7.08 nm/s at 0.8 μM; p-value: 0.40). **b**, At lower FtsA concentrations, the off-rate of FtsZ is similar to FtsA WT and R286W. At 0.8 μM FtsAs, FtsZ remains bound longer with FtsA R286W (0.20 ± 0.04 s^-1^ and 0.11 ± 0.01 s^-1^, p-value: 9.12*10^−4^). **c**, Single molecule lifetime of FtsZ is similar at 0.2 μM (7.31 ± 0.98 s and 8.31 ± 2.10 s, p-value: 0.58) and 0.4 μM FtsA (6.73 ± 1.23 s and 8.34 ± 1.53 s, p-value: 0.31). **d**, FtsA R286W recruits more FtsZ to the membrane than FtsA WT (p-value: 4.94*10^−5^). **e**, The persistency of the FtsZ pattern is lower with FtsA WT compared to FtsA R286W at all tested concentration, indicated by higher autocorrelation decay times τ. **f**, Scheme of TIRF experiment to measure the lifetime of a TAMRA labelled FtsZ C-terminal peptide. **g**, Representative histograms of the FtsZ CTP lifetime distribution in the presence of FtsA WT-His6 and FtsA R286W-His6. **h**, Representative curves for QCM-D experiments. At 0.8 μM the frequency change (=membrane binding) is identical between FtsA WT (green) and R286W (cyan). FtsA R286W can be washed easily from the membrane, whereas FtsA WT sticks stronger to the membrane, indicating stronger oligomerization. **i**, Scatter plots of FtsA wt (green, left) and FtsA R286W (cyan, right) intensities against FtsZ intensities from colocalization analysis for 0.2 and 0.4 μM. The steepness of the slope is consistently high for FtsA R286W but decreases drastically for FtsA WT at 0.4 μM.

**Figure S2:**
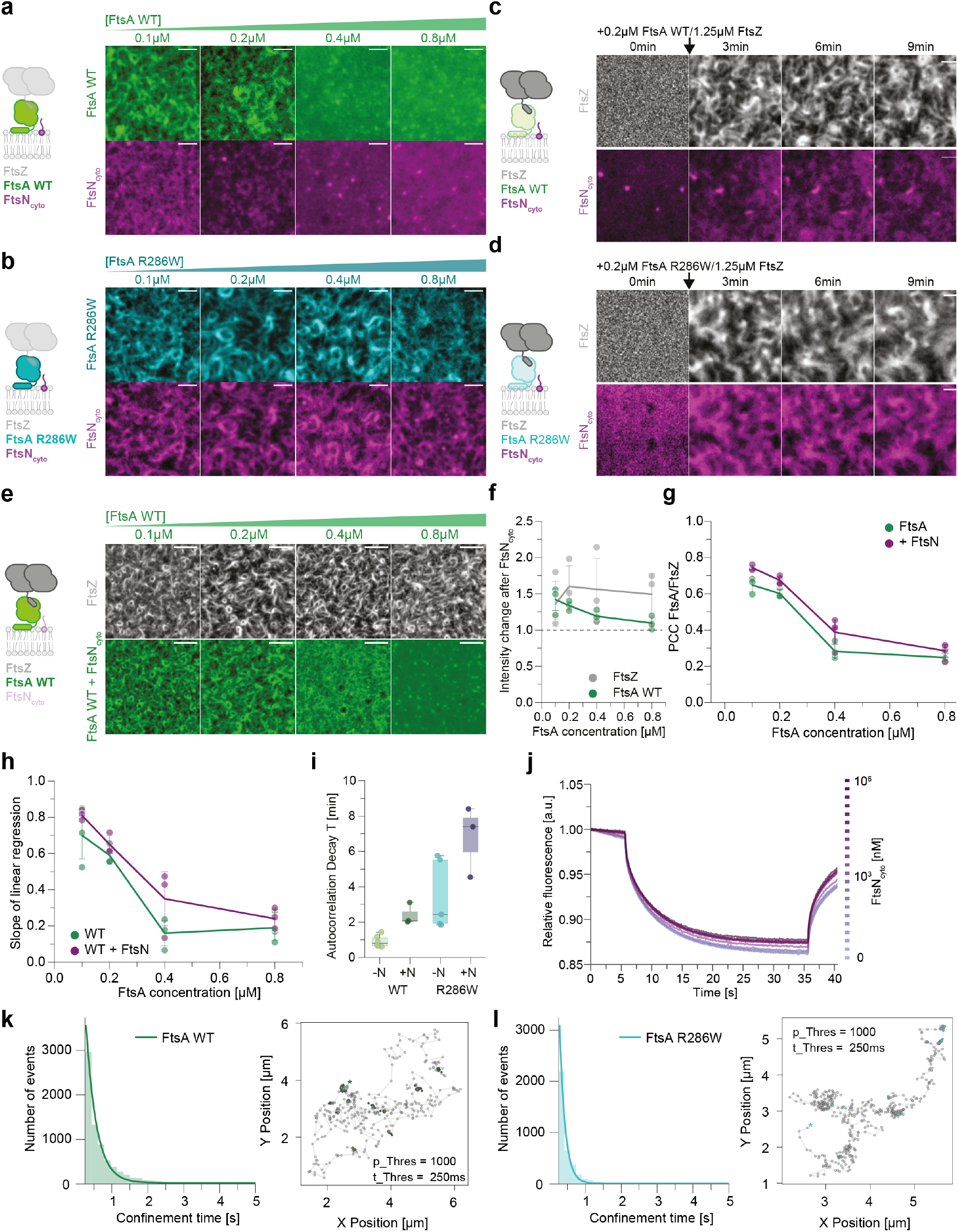
FtsN_cyto_ follows FtsA R286W co-filaments better than FtsA WT. **a**, Representative micrographs of Cy3-FtsA WT (green) and Cy5-FtsN_cyto_ (magenta) at increasing FtsA WT and constant FtsZ concentration and Tris-NTA-lipid ratio of 0.25%. FtsN_cyto_ and FtsA form co-filaments at FtsA concentrations of up to 0.2 μM, but fail to form a pattern at higher concentrations. **b**, Representative micrographs of Cy3-FtsA R286W (cyan) and Cy5-FtsN_cyto_ (magenta) at increasing FtsA R286W and constant FtsZ concentration. FtsN_cyto_ and FtsA form filaments in all concentrations tested. **c**, From a homogeneous distribution, Cy5-FtsN_cyto_ (magenta) colocalizes with FtsZ filaments on the membrane after the addition of A488-FtsZ (grey) and 0.2 μM FtsA WT at 0 min. **d**, From a homogeneous distribution, Cy5-FtsN_cyto_ (magenta) does colocalize with FtsZ filaments on the membrane after the addition of A488-FtsZ (grey) and 0.2 μM FtsA R286W at 0 min. Cartoons on the left side of **a-d** indicate the present and the labelled (bold) components. Scale bars are 2 μm. **e**, Representative micrographs of Cy3-FtsA WT (green) and A488-FtsZ (grey) presence of FtsN_cyto_. Scale bars are 5 μm. **f**, Presence of FtsN increases the amount of recruited FtsZ at all tested FtsA WT concentrations, whereas the amount of membrane bound FtsA remains constant above 0.2 μM. **g**, Colocalization of FtsA/FtsZ quantified by PCC shows that colocalization is slightly increased in the presence of FtsN_cyto_, but the effect is not significant (0.28 ± 0.04; 0.39 ± 0.07; p-value: 0.10; at 0.4 μM). **h**, The slope of the linear regression indicates the [FtsZ] vs. [FtsA] ratio. In the presence of FtsN, the slope for FtsA WT increases slightly. **i**, FtsN_cyto_ increases the persistency of the FtsZ pattern indicated by an increase in the autocorrelation decay time τ. **j**, Representative MST traces for a titration of FtsN_cyto_ to 50nM Cy5-FtsA WT. **k/l**, Histogram of FtsN_cyto_ confinement times to co-filaments of FtsZ and FtsA WT (**k**) or FtsA R286W (**l**). Representative tracks containing FtsN_cyto_ confinement events in the presence of FtsA WT (green) and R286W (cyan) are shown next to the histograms. The star indicates the beginning of the tracks. The insets indicate the chosen filters for the confinement analysis.

**Figure S3:**
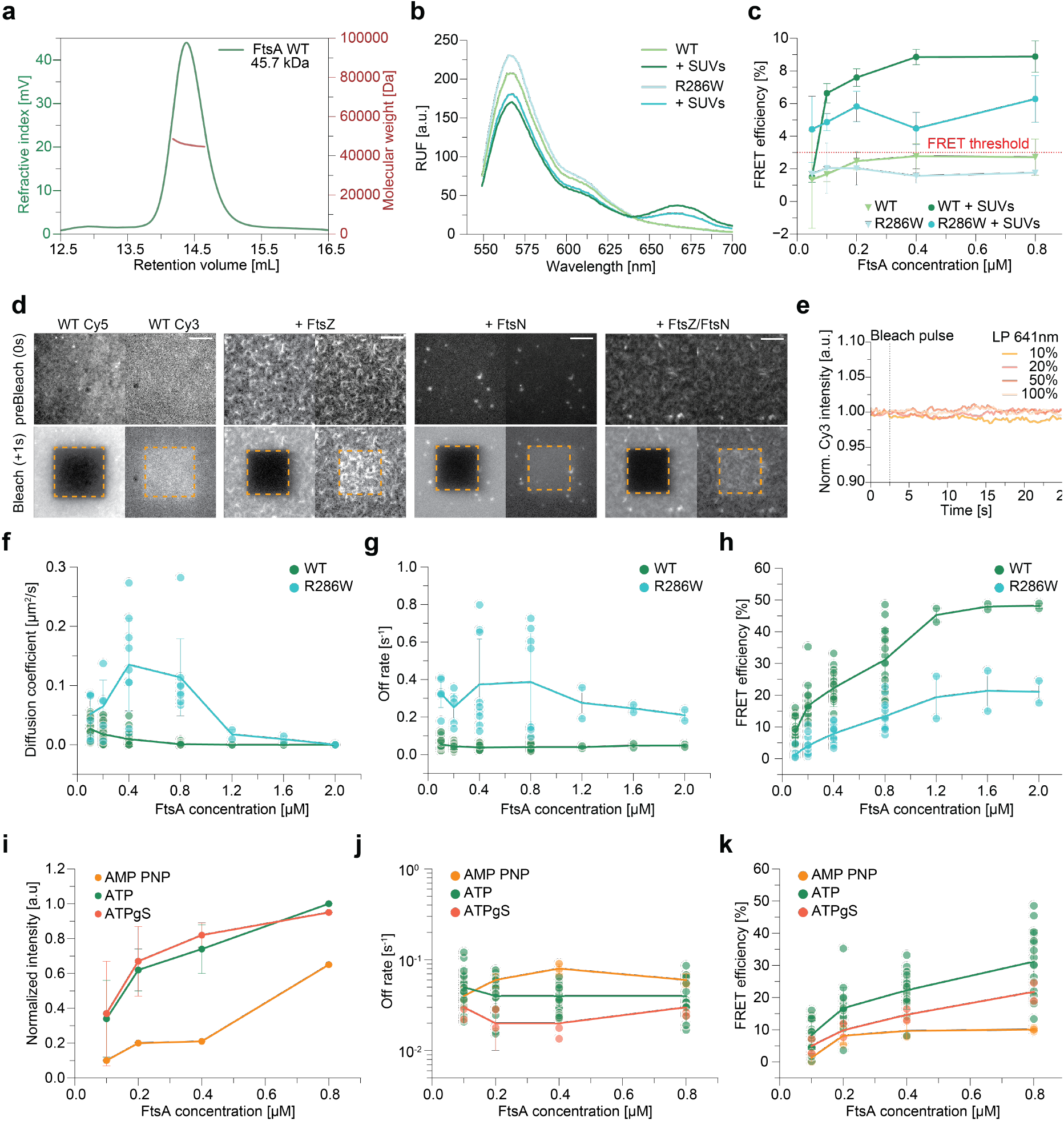
FtsA WT self-interaction is membrane dependent, but does not require ATP hydrolysis. **a**, SEC-MALLS experiments show that FtsA WT at 10 μM is a monomer in solution. **b**, Cuvette-FRET measurements indicate that self-interaction of both, FtsA WT (green) and FtsA R286W (cyan), depends on the presence of membranes. **c**, FRET signal is below the significance threshold at all concentrations tested in absence of vesicles. In the presence of lipids FtsA WT exhibits stronger FRET. **d**, Representative micrographs for acceptor photobleaching experiments of FtsA WT alone (top left), WT + FtsZ (top right), WT + FtsN (bottom left) and WT + FtsZ/FtsN (bottom right). Scale bars are 5 μm. **e**, The intensity of membrane-bound Cy3-FtsA WT is not affected by a Cy5-bleach pulse with increasing laser power (LP). **f**, Lateral diffusion of FtsA R286W drops significantly to FtsA WT levels at concentrations above 0.8 μM. The lateral mobility of R286W seems lower at low concentrations, as the fast off-binding rate dominates. Additionally, less protein is bound to the membrane as shown in QCM-D experiments, which impedes D_coeff_ analysis. **g**, The off-rate of FtsA R286W remains faster than for FtsA WT at all tested concentrations. **h**, FRET efficiency of FtsA R286W and FtsA WT saturates at concentrations above 0.8 μM. **i**, Nucleotide hydrolysis is not important for membrane binding of FtsA WT, as the protein binds to SLBs comparable in the presence of ATP and ATPγS. However, membrane binding is decreased in the presence of AMP PNP, which binds FtsA with lower affinity. **j**, The off-rate of FtsA WT is similar in the presence of ATP or ATPγS. **k**, FRET efficiency of FtsA WT is similar in the presence of ATP or ATPγS.

**Figure S4:**
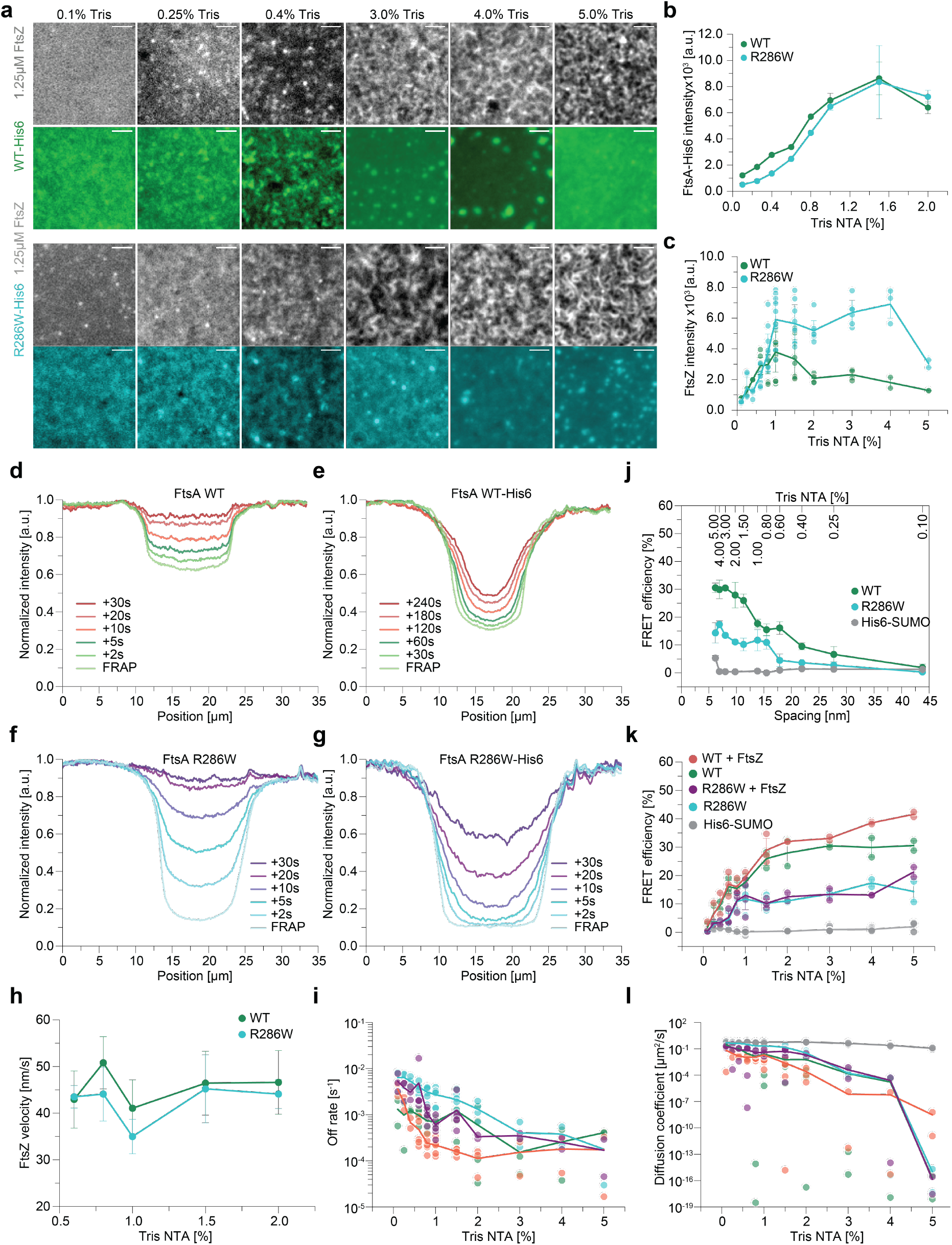
FtsZ pattern forms more efficient with FtsA R286W-His, even at high densities. **a**, By using His-tagged proteins and membranes with Tris-NTA lipids allow to control the density of FtsA on the membrane as demonstrated by the intensities of His-tagged FtsAs at different Tris-NTA lipid densities. **b**, At Tris-NTA lipid densities higher than 1%, FtsA R286W-His is able to recruit more FtsZ filaments than the FtsA-His. **c**, Representative micrographs of A488-FtsZ (grey) and Cy5-FtsA-His (green) or Cy5-FtsA R286W-His (cyan) at Tris-NTA lipid densities not shown in **Fig 4**. While the minimal protein density of FtsA needed to form an FtsZ pattern is similar, FtsZ filaments form a more disrupted cytoskeletal pattern at high densities of FtsA-His. **d-g**, Comparison of FRAP recovery profiles of native FtsAs **(d, f)** and His-tagged FtsAs **(e, g)**. The recovery of the native proteins is dominated by exchange, while His-tagged proteins recovers dominantly by lateral diffusion, as can be seen by the different shapes of the recovery profiles. **h**, The treadmilling speed of FtsZ is similar at different densities of FtsA-His or FtsA R286W-His. **i**, As expected, off-rates of FtsA-His and FtsA R286W-His obtained from FRAP experiments are very low. **j**, FRET of His-tagged FtsAs decreasing with increasing spacing indicating self-interaction. The His6-SUMO control only exhibits weak FRET at the maximum density were different fluorophores can be closer to each other than the theoretical FRET limit (5 nM). **k**, FRET efficiency of His-tagged FtsAs ± FtsZ and His6-SUMO on membranes with up to 5% Tris-NTA lipids. The FRET signal of FtsA-His is consistently higher compared to FtsA R286W-His. Adding FtsZ has a only a very modest effect on the self-interaction. **l**, Diffusion coefficient of His-tagged FtsAs ± FtsZ and His6-SUMO on membranes with up to 5% Tris-NTA lipids. While mobility of His6-SUMO does not change, His-tagged FtsAs diffuses slower with increasing densities. Addition of FtsZ further decreases diffusion.

## Methods

### Reagents

All of the reagents, chemicals, peptides and software used are listed in the reagents table.

### Purification and fluorescence labelling of FtsZ

FtsZ was purified as previously described^4,21^. In short FtsZ with a -terminal His_6_-SUMO fusion protein and seven residues (AEGCGEL) for maleimide coupling of thiol-reactive dyes was cloned into a pTB146-derived vector^21^. FtsZ was expressed in *E. coli* BL21 cells, at 37 °C in Terrific Broth supplemented with 100 μg ml^−1^ ampicillin and expression was induced at an OD600 of 0.6-0.8 with 1 mM isopropyl-β-thiogalactopyranoside (IPTG) and incubated for 5 h at 37 °C. Cells were harvested by centrifugation (5,000*g* for 30 min at 4 °C). The pellet was resuspended in buffer A (50 mM Tris-HCl [pH 8.0], 500 mM KCl, 2 mM β-mercaptoethanol and 10% glycerol) plus 20 mM imidazole and supplemented with ethylenediaminetetraacetic acid (EDTA)-free protease inhibitor cocktail tablets (Roche Diagnostics). Cells were lysed using a cell disrupter (Constant Systems; Cell TS 1.1) at a pressure of 1.36 kbar and subsequently incubated with 2.5 mM MgCl_2_ and 1 mg ml^−1^ DNase for 15 min. Cell debris was removed by centrifugation at 60,000*g* for 30 min at 4 °C and the supernatant was incubated with nickel-nitrilotriacetic acid (Ni-NTA) resin (HisPur Ni-NTA Resin; Thermo Fisher Scientific) for 1 h at 4 °C. The resin was washed with buffer A containing 10 mM imidazole, followed by buffer A with 20 mM imidazole and the protein was subsequently eluted with buffer A supplemented with 250 mM imidazole. To cleave the His6-SUMO, FtsZ together with His-tagged SUMO protease (Ulp1) (1:100 molar ratio) was dialyzed overnight at 4 °C against buffer B (50 mM Tris-HCl [pH 8.0], 300 mM KCl and 10% glycerol). To remove remaining His-tagged molecules, the sample was again passed through Ni-NTA resin, equilibrated with buffer B. The polymerization-competent fraction of the purified FtsZ was enriched by CaCl_2_ at room temperature after buffer exchange into polymerization in buffer C (50 mM PIPES [pH 6.7] and 10 mM MgCl_2_. Polymerization was induced with 10mM CaCl_2_ and 5mM GTP, incubated for 20 mins at RT and the polymeric fraction was collected by centrifugation at 15,000*g* for 2 min and the gel-like pellet was resuspended in buffer D (50 mM Tris-HCl [pH 7.4], 50 mM KCl, 1 mM EDTA and 10% glycerol). For labelling, the thiol-reactive dye Alexa Fluor 488 C5 Maleimide (Thermo Fisher Scientific) was dissolved in dimethyl sulfoxide (DMSO) following the manufacturer’s instructions. FtsZ was reduced by incubating the protein with a 100× molar excess of tris(2-carboxyethyl)phosphine (TCEP) for 20 min at room temperature. A 10× molar excess of Alexa Fluor 488 was added and extensively dialyzed against buffer D overnight at 4 °C. Remaining CaCl_2_, GTP and free dye was removed via a PD10 desalting column and peak fraction were collected, flash frozen in liquid nitrogen and stored at −80 °C.

### Purification and fluorescence labelling of FtsAs

FtsA was cloned into vector pMAR19, with an N-terminal TwinStrep-SUMO fusion protein plus a 5xGlycine tag for fluorescence labelling via sortagging. FtsA was expressed in *E. coli* BL21 cells, grown at 37 °C in 2× YT medium supplemented with 100 μg ml^−1^ ampicillin and expression was induced at an OD600 of 0.6-0.8 with 1 mM IPTG. The protein was expressed overnight at 18 °C and harvested by centrifugation (5,000*g* for 30 min at 4 °C. The pellet was resuspended in buffer A (50 mM Tris-HCl [pH 8.0], 500 mM KCl, 10 mM MgCl_2_ and 0.5mM DTT) supplemented with EDTA-free protease inhibitor cocktail tablets and 1 mg ml^−1^ DNase I. Cells were lysed by sonication using a Q700 Sonicator equipped with a probe of 12.7mm diameter, which was immersed into the resuspended pellet. The suspension was kept on ice during sonication (Amplitude 40, 1sec on, 5sec off for a total time of 10 minutes). Subsequently, cell debris was removed by centrifugation at 23,500*g* for 45 min at 4 °C. The clarified lysate was incubated with IBA Lifesciences Strep-Tactin® Sepharose® resin for 1h at 4 °C. Subsequently, the resin was washed with 40x CV buffer A and the fusion protein was eluted using buffer A containing 5 mM desthiobiotin. The protein concentration was determined with Bradford and adjusted to 12μM with buffer A, in order to avoid precipitation of the protein. The His6-SUMO protease Ulp1 was added in a 1:100 molar ratio and the TS-SUMO tag was cleaved overnight at 4°C, without shaking. To remove the cleaved tag and Ulp1, FtsA was subjected so Size Exclusion Chromatography. A HiLoad 26/600 Superdex 200 Prep grade column was equilibrated with buffer B (50mM Tris [pH 8.0], 500mM KCl, 10mM MgCl_2_, 10% Glycerol and 0.5mM DTT) and the protein was injected. The peaks containing the final protein, corresponding to monomeric FtsA, were determined via SDS-page gel-electrophoresis, pooled and concentrated as described above. For total internal reflection fluorescence microscopy FtsA was labeled with Cyanin-3 or Cyanin-5 via sortagging^40^. The 5xGly tag at the N-terminus of FtsA was conjugated to CLEPTGG-peptide, which was previously labeled via maleimide directed labelling with either sulfo-Cyanine 3 or Cyanine 5-maleimide (Lumiprobe). 10μM Sortase, 0.5mM labeled peptide and 10μM of FtsA were mixed together and incubated overnight at 4°C. To remove free peptide, free dye and sortase, FtsA was subjected to another Size-exclusion on a HiLoad Superdex 200 16/600 prep grade column, pre-equilibrated with buffer B. The monomeric protein was collected and the concentration was determined via Bradford. The concentration of dye molecules (=labeled protein) was measured by NanoDrop and the Degree of Labelling (DoL) was determined by calculating the ratio of Labeled Protein:Protein. The DoL for FtsAs was between 65-70%. To obtain the hypermorphic mutant of FtsA, R286W, pMAR19 was used as a base for site-directed mutagenesis (SDM). We replaced Arginine 286 with Tryptophan, by exchanging a single nucleotide (C → T), resulting in pMAR25. The variant of FtsA was purified in the same way as described above for the wildtype protein.

To purify His-tagged variants of FtsA wt and R286W, the C-terminal amphipathic helix at position 405-420 (GSWIKRLNSWLRKEF*) of pMAR19/pMAR25 was replaced by a 6xHistidine Tag, resulting in pNB4 and pNB5 (FtsA WT-His6 and R286W-His6 respectively). The purification and labeling were performed as described above for native FtsA wt.

### Purification and fluorescence labelling of His-tagged SUMO-Cys (HS-Cys)

As a control for the FRET assay, we constructed a vector based on pTB146 containing only the SUMO protein, modified with a N-terminal 6xHis tag and a C-terminal Cysteine for maleimide labelling, resulting in pPR5. HS-Cys was expressed in *E. coli* BL21 cells, at 37 °C in Terrific Broth supplemented with 100 μg ml^−1^ ampicillin and expression was induced at an OD600 of 0.6-0.8 with 1 mM isopropyl-β-thiogalactopyranoside (IPTG) and incubated for 3 h at 37 °C. Cells were harvested by centrifugation (5,000*g* for 30 min at 4 °C). Lysis and incubation with Ni-NTA-beads was performed as described before for FtsZ. The protein was eluted with buffer A (50 mM Tris-HCl [pH 7.4], 300 mM KCl and 10% glycerol) supplemented with increasing concentrations of Imidazole (50/100/150/200/250/300/400mM). The purity of the fractions were checked by SDS Page and the eluted fractions with 200 and 250mM Imidazole were pooled together. Subsequently HS-Cys was dialyzed overnight against buffer B (50 mM Tris-HCl [pH 7.4], 100 mM KCl and 10% glycerol) to remove remaining Imidazole. HS-Cys was labelled by maleimide directed labelling with sulfo-Cyanine 3 and Cyanine-5 maleimide as described above for FtsZ. To remove free dye and remaining traces of Imidazole, the protein was subjected to Size-exclusion on a HiLoad Superdex 200 16/600 prep grade column, pre-equilibrated with buffer B. The peaks, corresponding to the labelled HS-Cys were collected and concentrated with Vivaspin 20 centrifugal concentrators (5kDa cutoff). The final concentration and the degree of labeling were determined as described above and the protein was stored at -80°.

### Labeling of peptides

The cytoplasmic peptide of FtsN with a C-terminal His6 tag and an N-terminal cysteine residue was labelled and handled as described before^4^. The C-terminal peptide (CTP) of FtsZ conjugated to an N-terminal TAMRA dye (5-Carboxytetramethylrhodamine) (TAMRA-KEPDYLDIPAFLRKQAD) was purchased from Biomatik and reconstituted in buffer A (50mM HEPES-KOH, [pH 7.4] to a concentration of 2mg ml^-1^, flash frozen and stored at −80 °C.

### Quartz crystal microbalance-Dissipation (QCM-D)

QCM-D experiments were performed with the QSense Analyzer from Biolin Scientific, equipped with silica coated sensor (QSX 303). The sensors were cleaned for 10s in a Zepto plasma cleaner, mounted in the QCM-D chambers and the acquisition of the experiment was first performed in the reaction buffer (50mM Tris-HCl [pH 7.4], 150mM KCl and 5mM MgCl_2_. The supported lipid membrane was formed by rupturing 0.5mM small unilamellar vesicles (SUVs) in the presence of MgCl_2_. The lipid composition used was 67% DOPC : 33% DOPG. After bilayer formations, the reaction buffer supplemented with 2mM ATP, 2mM GTP and 1mM DTT was injected and the signal was recorded until a stable equilibrium was reached. Subsequently increasing concentrations of FtsA were injected in the QCM-D chamber and changes in frequency and dissipation were monitored in real-time. The flow rate used in all experiments was 25μL min^-1^ and the temperature was set to 25 °C. To estimate the membrane binding affinity, we extracted the frequency changes for different FtsA concentrations and fitted a Hill equation 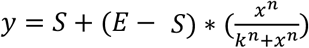, where S is the starting point, E the end point, n is the Hill coefficient and k the dissociation constant. The measured binding affinity is an upper estimate, because QCM-D accounts not only on the dry molecular mass, but also on hydrational shell of the molecular assembly.

### Size exclusion chromatography with multiple angle light scattering (SEC-MALS)

45μg (100μL of a 12μM solution) of purified FtsA WT was resolved on a Superdex 200 Increase 10/300 at a flow rate of 0.5ml min^-1^ at room temperature. Light scattering was recorded on a miniDawn light scattering device (Wyatt). Changes in the refractive index were used to define the peak area, which was used to obtain the molecular mass. The analysis of the data was performed with the ASTRA software (Wyatt).

### Microscale thermophoresis (MST)

MST experiments were performed with either 50 nM FtsA wt or R286W labeled with Cyanin-5 and increasing concentrations of unlabeled FtsN_cyto_. The peptide was diluted in buffer A (50 mM Tris-HCl [pH 7.4], 150 mM KCl, 5 mM MgCl_2_ and 0.005% Tween-20). After adding the peptide to the protein, the mixtures were left to incubate for 10 min at room temperature and subsequently loaded in premium coated capillary tubes (NanoTemper). Measurements were performed with a Monolith NT.115 (NanoTemper) equipped with a blue and a red filter set. The data was acquired with 20% MST and 20% light-emitting diode settings at 25ºC. Cy5 fluorescence was measured for 5s before applying a thermal gradient for 30 s. Binding curves were obtained by plotting the normalized change in fluorescence intensity after 20s against the concentration of titrated peptide. To extract the binding affinity, a Hill equation was 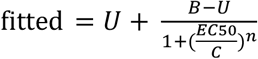, where C is the peptide concentration, U is the signal for the unbound state, B the signal for the bound state and n is the Hill coefficient.

### Cuvette FRET experiments

For in solution FRET experiments, increasing concentrations of Cy3-FtsA and Cy5-FtsA in a 50%:50% ratio were mixed in 100μL reaction buffer (50mM Tris-HCl [pH7.4], 150mM KCl and 5mM MgCl2) inside a quartz cuvette (Hellma® fluorescence cuvettes, ultra Micro). The spectrums were measured using a Spectrophotometer Spectramax M2e Plate-+ Cuvette Reader. Cy3 labeled FtsA was excited at a wavelength of 520nm and the resulting emission spectrum was recorded from 550-700nm in 1nm steps. To avoid crosstalk of the excitation light, a cutoff filter was set to 550nm. Addition of ATP or small unilamellar vesicles (SUVs) was measured individually for each concentration. Buffer controls, containing the corresponding reagents and only Cy5-FtsA were measured and used as background corrections for measurements Cy3-& Cy5-FtsA. Background corrected spectra were used to estimate FRET efficiency by 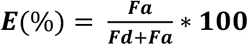, where Fa is the peak of acceptor (=Cy5) emission at 670nm and Fd the peak of the emission of the donor (=Cy3) spectrum at 565nm.

### Preparation of coverslips

Glass coverslips were cleaned in piranha solution (30% H_2_O_2_ mixed with concentrated H_2_SO_4_ at a 1:3 ratio) for 60 min, and extensively washed with ddH_2_O, followed by 10min sonication in double-distilled H_2_O and further washing with ddH_2_O. Cleaned coverslips were stored for no longer than 1 week in H_2_O water. Before formation of the supported lipid bilayers, the coverslips were dried with compressed air and treated for 10 min using a Zepto plasma cleaner (Diener electronics) at full power. As reaction chambers, 0.5-ml Eppendorf tubes, without the conical end, were glued on the coverslips with ultraviolet glue (Norland Optical Adhesive 63) and exposed to ultraviolet light for 10 min.

### Preparation of small unilamellar vesicles (SUVs)

For experiments without His tagged peptides, 1,2-dioleoyl-sn-glycero-3-phospho-(1’-rac-glycerol) (DOPC) and 1,2-dioleoyl-sn-glycero-3-phospho-(1^′^-rac-glycerol) (DOPG) at a ratio of 67:33 mol% was used. To enable peptide attachment to the lipid membrane SUVs with 1 mol% dioctadecylamine (DODA)-tris-NTA (synthesized by ApexMolecular), in a ratio of 66:33:1 mol% DOPC:DOPG:Tris-NTA, were prepared. To titrate the density of Tris-NTA lipids, SUVs without and with up to 5% Tris-NTA were mixed together before supported lipid bilayer formation in the appropriate volumes. For SUV preparation, lipids in chloroform solution were added into a glass vial and dried with filtered N_2_ to obtain a thin homogeneous lipid film. Residual chloroform was removed by further drying the lipids for 2-3h under vacuum. Subsequently swelling buffer (50 mM Tris-HCl [pH 7.4] and 300 mM KCl) was added to the lipid film and incubated for 30 min at room temperature to obtain a total lipid concentration of 5 mM. When Tris-NTA lipids were present in the mix, 5mM Ni_2_SO_4_ were added to the swelling buffer to load NTA groups with Nickel. To disrupt multilamellar vesicles, the mixture was repeatedly vortexed rigorously and freeze– thawed (8x) in dry ice or liquid N_2._ To obtain small unilamellar vesicles the liposome mixture was tip-sonicated using a Q700 Sonicator equipped with a 1/2mm tip (amplitude =1, 1s on, 4s off) for 25 min on ice. The vesicles were centrifuged for 5 min at 10,000*g* and the supernatant was stored at 4 °C in an Argon atmosphere and used within 1 week.

### Preparation of supported lipid bilayers (SLBs)

To prepare supported lipid bilayers, the SUV suspension was diluted to a lipid concentration of 0.5 mM with swelling buffer. Vesicle rupture was induced by adding 5mM CaCl_2_ to the SUVs on the glass surface. The bilayers were incubated for 30 minutes at 37ºC, and remaining non-fused vesicles were washed away by pipetting an excess of swelling buffer (5x) on top, followed by 5x washes with reaction buffer (50 mM Tris-HCl [pH 7.4], 150 mM KCl and 5 mM MgCl_2_) The membranes were used within 4 hours after the preparation.

### TIRF microscopy

Experiments were performed using two TIRF microscopes. The iMIC TILL Photonics was equipped with a 100× Olympus TIRF NA 1.49 differential interference contrast objective. The fluorophores were excited using laser lines at 488, 561 and 640 nm. The emitted fluorescence from the sample was filtered using an Andromeda quad-band bandpass filter (FF01-446-523-600-677). For the dual-colour experiments, an Andor TuCam beam splitter equipped with a spectral long pass of 580 and 640 nm and a band pass filter of 525/50, 560/25 and 710/80 (Semrock) was used. Time series were recorded using iXon Ultra 897 EMCCD Andor cameras (X-8499 and X-8533) operating at a frequency of 5 Hz for standard acquisition and at 10 Hz for single-molecule tracking. The Visitron iLAS2 TIRF microscope was equipped with a 100xOlympus TIRF NA 1.46 oil objective. The fluorophores were excited using laser lines at 488, 561 and 640 nm. The emitted fluorescence from the sample was filtered using a Laser Quad Band Filter (405/488/561/640 nm). For the dual-colour experiments, a Cairn TwinCam camera splitter equipped with a spectral long pass of 565 and 635 nm and band pass filters of 525/50, 595/50, 630/75, 670/50 and 690/50 was used. Time series were recorded using Photometrics Evolve 512 EMCCD (512 × 512 pixels, 16 × 16 μm^2^) operating at a frequency of 5 Hz for standard.

### Dual color FtsA-FtsZ experiments

To study co-localization and co-treadmilling of FtsA with treadmilling FtsZ filaments on supported lipid bilayers, we used Cy5-FtsA wt or Cy5-R286W (0.1–0.8 μM) and FtsZ-A488 (1.25μM) in 100 μl of reaction buffer. Additionally, the reaction chamber contained 4 mM ATP and 4 mM GTP, as well as a scavenging system to minimize photobleaching effects: 30 mM d-glucose, 0.050 mg ml−1 Glucose Oxidase, 0.016 mg ml−1 Catalase, 1-10 mM DTT and 1 mM Trolox. Prior addition of all components a corresponding buffer volume was removed from the chamber to obtain a total reaction volume of 100 μl. The dynamic protein pattern was monitored by time-lapse TIRF microscopy at one frame per two seconds and 50-ms exposure time.

### FtsZ single molecule experiments

Single molecule experiments were performed as described previously^41^. In short, individual FtsZ proteins were imaged at single molecule level by adding small amounts of Cy5-labelled FtsZ (200pM) to a chamber with 0.2/0.4 μM FtsA wt/R286W and 1.25 μM A488-FtsZ.

### Single molecule measurements of the C-terminal peptide of FtsZ (CTP)

To measure residence times of the FtsZ-CTP, we added the TAMRA-labeled CTP peptide (TAMRA-KEPDYLDIPAFLRKQAD, synthesized by Biomatik) to membranes with 1% Tris-NTA lipids. Before addition of the peptide, 1μM of His-tagged variants of FtsA wt and R286W were added to the chambers, incubated for 20 minutes and washed 6x with reaction buffer. Subsequently, 1nM of FtsZ TAMRA-CTP was added to the chamber and single molecule time-lapses were acquired every 32 or 51 ms, with exposure times of 30 and 50 ms, respectively.

### Dual-color FtsN-FtsZ and FtsN-FtsA experiments

To study the colocalization of Cy5-labeled FtsN_cyto_, we used membranes with 0.25% Tris-NTA lipids to ensure stable peptide immobilization. FtsN peptide at the concentration of 1μM was added to the chamber and left to incubate for 20 minutes to ensure homogeneous binding. Subsequently the chamber was washed 6x with reaction buffer, to remove bulk peptide. To visualize colocalization with either FtsZ or FtsA, either a mix of FtsA and FtsZ-A488 or Cy3-FtsA and FtsZ was added. The concentration of FtsZ was again kept constant at 1.25μM, whereas FtsA concentrations were titrated from 0.1-0.8 μM. The time-lapse videos were recorded for 10 minutes after the addition, with one frame per two seconds.

### Single molecule experiments for confinement of FtsN_cyto_

To study the interaction of single molecules of Cy5-labeled FtsN_cyto_, we also used membranes with 0.25% Tris-NTA lipids. This time 1μM of unlabeled FtsN_cyto_ supplemented with 50 pM of Cy5-labeled FtsN_cyto_ were added to the chamber and incubated for 20 minutes, followed by 6x washes with reaction buffer. Subsequently, 0.2 μM of either FtsA WT or FtsA R286W and 1.25 μM FtsZ-A488 were added to the chamber and pattern formation was recorded for 10 minutes. Single molecule time-lapses were acquired every 32 or 51 ms, with exposure times of 30 and 50 ms, respectively.

### Single molecule experiments on FtsA WT and FtsA R286W

To study the behaviour of single molecules of FtsA WT and FtsA R286W, 0.1μM of the unlabeled protein supplemented with 35pM Cy5 of the respective FtsA variant were added to the reaction chamber. After 5 minutes of incubation, single molecule time-lapses were acquired every 125, 250, 500, and 1000 and 2000 ms, with an exposure time of 50 ms. Subsequently, the bulk concentration of FtsA was increased to 0.2/0.4/0.8 μM and single molecule time-lapses were repeated as described above.

### FRAP and FRET experiments on SLBs

To measure the membrane residence time and self-interaction of FtsA WT and FtsA R286W, acceptor (Cy5) photobleaching experiments were performed. Equimolar concentrations of Cy3- and Cy5-labeled FtsA, supplemented with 20% unlabeled FtsA were used to study FRET and FRAP. Five pre-bleach frames were acquired, followed by acceptor photobleaching of a rectangular ROI with 40% 641 laser power and a dwell size of 1μs/pixel, 75% overlapping lines. The recovery of the signal or the increase in donor intensity were measured with either 2 frames or 1 frame per second. The different acquisition rates were implemented, due to the accelerated recovery of R286W compared to wt. To measure effects of FtsZ, 1.25 μM of unlabeled FtsZ were added to the membrane and FRET/FRAP was measured again. To quantify effects of FtsN, SLBs with 0.25% Tris NTA-lipids were pre-equilibrated with FtsN_cyto_ before additions of FtsAs. Subsequently, 1.25 μM FtsZ was added as well to quantify effects of the combined presence of FtsN and FtsZ.

### SLB experiments of His-tagged FtsAs

To study colocalization of His-tagged variants of FtsA, 0.5-1μM of Cy5 labeled His-tagged FtsAs were added to the chamber, incubated for 20 minutes and washed 6x with reaction buffer. Subsequently 1.25 μM A488-FtsZ was added to the chamber and pattern formation was recorded for 20 minutes. To control the density of membrane bound FtsA, SUVs without and with Tris-NTA lipids were mixed to obtain the respective Tris-NTA concentrations. To perform FRET/FRAP experiments, equimolar concentrations of Cy3 & Cy5 labeled His-tagged FtsAs (total 0.5-1 μM) or His-SUMO-Cys were added to a chamber and treated as above. To study effects of FtsZ, 1.25 μM unlabeled FtsZ were added to the chamber and recorded for 20 minutes. FRAP experiments were performed as described before.

### Image processing and analysis

For data analysis, the movies were imported to the FIJI software^42^. For data analysis, raw, unprocessed time-lapse videos were used. All micrographs in the manuscript were processed with the walking average plugin of ImageJ, averaging the signal of four consecutive frames, and contrast was optimized for best quality.

### Colocalization analysis

Time-lapse videos were first intensity-corrected and contrast-enhanced to avoid bleaching effects and simplify subsequent analysis. To remove contributions of X-Y drift, the videos were processed with the Linear Stack Alignment with SIFT plugin. Proper alignment was checked with the 3TP align plugin (J. A. Parker; Beth Israel Deaconess Medical Center, Boston). Subsequently, regions of interest (ROIs) in the center of the stacks were chosen for colocalization analysis. The Pearson’s correlation coefficient (PCC) was quantified with the Image CorrelationJ 1o plugin. To extract information about the relative ratio of FtsZ/FtsA molecules (slope of linear regression) we also used the Image CorrelationJ 1o plugin. As an output, the plugin provides a scatterplot of FtsZ vs. FtsA intensities, to which we fitted a linear slope *y*=*k* * *x* + *d*, where k is the slope and d the offset. The slope was used as an estimate for the ratio of FtsA molecules below FtsZ filaments^43^.

### FtsN recruitment rate quantification

To estimate the rate of FtsN_cyto_ recruitment towards FtsA/Z co-filaments, we measured the PCC after adding FtsA/Z to a membrane homogeneously covered with FtsN_cyto_ and fitted a power law equation *y*=*a \** (1− *e*^*−b*t*^) + *c*, where a is the starting point, b is the rate and c is the offset, to the increasing PCC values after protein addition and extracted the recruitment rate.

### Treadmilling and temporal PCC analysis

Treadmilling dynamics, were quantified using an automated image analysis protocol previously developed in our lab^25^. To visualize colocalization of the co-treadmilling FtsZ & FtsA filaments, we used dual-color videos obtained at an acquisition rate of one frame per two seconds. The two channels were aligned using FIJI’s 3TP align plugin. Both channels were then subjected to the image subtraction protocol and colocalization was measured as described above.

### FtsZ autocorrelation analysis

To measure reorganization dynamics of FtsZ filaments, we used a temporal correlation analysis based on the Image CorrelationJ 1o plugin. We quantified the PCC between the first frame to subsequent frames with increasing time lag (Δt). The decrease in the PCC was plotted against Δt to obtain autocorrelation curves. Slower decay indicates more persistent structures. The rates of decay were extracted by fitting monoexponential decay to the autocorrelation curves *y*=*a * e*^(*−b*t*)^ + *k*, where a is the starting point, b is the decay rate and k is the final offset. The half time of the monoexponential decay was calculated via the decay rate.

### Transient Confinement Analysis of FtsN_cyto_

Single molecule experiments with FtsN_cyto_were tracked using the TrackMate plugin from ImageJ^44^. Non-moving particles and short tracks (below 1s) were filtered out and the data exported as .xml files. To analyze transient confinement periods of FtsN_cyto_ to FtsZ/FtsA co-filaments we used the packing coefficient (*p*) to identify when diffusing FtsN_cyto_ molecules switch between free diffusion and confined motion. The packing coefficient is defined as the length of the trajectory in a short time window and the surface area that it occupies. This gives an estimate of the degree of free movement that a molecule displays in a period independently of its global diffusivity. This approach is adapted from Renner et al and implemented here as an easy-to-use python script^45^. The packing coefficient is computed for each time point as:

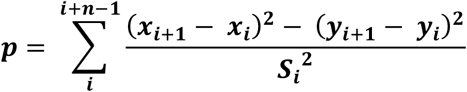

Where *xi,yi* are the coordinates at time *i, xi+1, yi+1* are the coordinates at time *i+1*, n is the length of the time window, and *Si* is the surface area of the convex hull of the trajectory segment between time points *i* and *i+n*. Periods of confinement are identified by setting a threshold corresponding to a certain confinement area size, since *p* scales with the size of the confinement area. Then it is possible to calculate the frequency and duration of confinement periods and to localize them in space. Each position will have a characteristic *p*, considering the behaviour of the following *n* positions. This approach overcomes the limitations of using MSD calculation, which overlooks transient confinement periods. Nevertheless, the Brownian diffusion trajectories can temporarily mimic confinement due to random fluctuations of the length of the displacements. However, the amplitudes and durations of these fluctuations are most of the time smaller and shorter than the ones associated with real non-Brownian transient motion. Therefore, the use of a threshold value of p (*P*_*thresh*_) and a minimal duration above this threshold (*t*_*thresh*_) can suppress the detection of apparent non-random behaviours without excluding the detection of real confinement. These parameters depend on the acquisition frequency (which will affect the length of the time window) and the characteristic time of confinement. Too large windows will not detect properly the confinement period, while the statistical uncertainty increases in shorter windows. Thus, to accurately detect confinement periods, the window size should be adjusted accordingly to the acquisition rate. To detect periods of confinement, we set *p*_*thres*_ to 1000, which corresponds to confinement areas of roughly <50nm, and a *t*_*thres*_ of 0.25 seconds, which corresponds to 5 and 8 frames when using 51msec and 32msec acquisition rates, respectively. The thresholds for confinement and time were chosen after manual inspection of tracks and corresponding confinement events. Finally, mean confinement times were extracted by fitting a monoexponential decay function to histograms of confinement times of individual experiments. To validate the performance of our code, we simulated single molecule tracks that switch between free diffusion and transient confinement periods using FluoSim^46^. To simulate appropriate tracks some parameters were kept constant for all tracks: 50 molecules/FOV; a D_coeff_ out- and inside 0.2μm^2^/s and the crossing probability was set to 1. To test the performance of our code we varied binding rates from 0.1-0.5s^-1^, unbinding rates from 0.5-3s^-1^ and the trapped D_coeff_ from 0.002-0.008μm^2^/s. The values were chosen according to previously acquired and published data^4^. At low diffusion coefficient for trapped molecules (<0.005 μm^2^/s) confinement periods were identified with a marginal error of ±0.04s, whereas the performance of the routine suffered slightly when increasing the diffusion coefficient of trapped molecules (> 0.008 μm^2^/s), but still resulted in values close to the ground truth (±0.1s). We also used these simulated tracks to fine-tune the thresholds for FtsN confinement time analysis. The source code can be found in https://github.com/paulocaldas/Transient-Confinement-Analysis.

### Single-molecule analysis of FtsZ and FtsA

Single molecules of FtsA or FtsZ were tracked using the TrackMate plugin from ImageJ^44^. To obtain the residence time of FtsZ and FtsA, we performed a residence time analysis as described before^41^. Shortly, single molecules were imaged at different acquisition rates (0.1-2s) and the lifetime of the molecules was extracted from each data set. To account for photobleaching effect, the obtained lifetimes were plotted against the acquisition rate. Then, we fitted a linear regression to this data and the photobleach corrected lifetime was calculated by taking the inverse of the slope of the linear regression. Furthermore, we extracted the diffusion coefficient of FtsA single molecules at increasing concentrations. For this we filtered the obtained data, by considering only trajectories which are present on the membrane for more than 0.4s. This low filter threshold was necessary, due to the very short lifetime of FtsA R286W molecules at low concentrations. Subsequently the diffusion coefficient of FtsA molecules was estimated by fitting an MSD curve to each individual trajectory.

### Quantifying FRET, D_coeff_ and k_off_ from FRAP experiments

To estimate the degree of self-interaction of FtsA WT and FtsA R286W, we used a photobleaching approach as outlined above. Bleaching the acceptor dye leads to an increase in the donor intensity, which can be used to quantify the Foerster Resonance energy transfer (FRET)^47^. FRET efficiency was quantified with 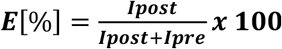, where I_post_ is the intensity of the acceptor (Cy3) after bleaching of the donor and I_pre_ is the acceptor intensity before bleaching the donor. While quantifying membrane binding dynamics of FtsA, we realized soon, that the recovery of FtsA was achieved by two different mechanisms: simple on- and off-binding to the membrane and lateral diffusion of the protein along the SLB. To extract the contribution of both processes, we adjusted a routine recently published by Gerganova *et al*.^35^. In short, the code provides the contribution of both modes of recovery by analyzing the shape of the fluorescence recovery profile. For simple on/off binding, the profile shape during recovery does not change (compare Fig. S4h left). Contribution of diffusion leads to a change of the slope of the outer borders of the bleached region (S4h right). Thus, by measuring the change of the slope of the border recovery, diffusion and simple on/off binding can be distinguished. To adjust the code to our needs, we created a wrapper around your fit function, which can be used directly on .tif images with support for ImageJ ROIs. We added a bleach correction, choice of projection axis (x or y) and optional mirroring, if bleaching was not symmetric. The original code by David Rutkowski can be found in https://github.com/davidmrutkowski/1DReflectingDiffusion, whereas our adjusted version, termed “FRAPdiff”, can be found at https://git.ist.ac.at/csommer/frapdiff.

### Calculation of spacing

To calculate the theoretical spacing of His-tagged FtsAs or the His SUMO control, we consulted a previous QCM-D study using Tris-NTA lipids and a His-tagged version of ZipA^48^. The spacing in nm^2^ was calculated by 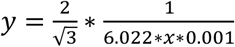, where x is the protein density in pMol cm^-2^ which can be estimated from the Tris-NTA lipid density^49^.

### Statistics and reproducibility

Statistical details of the experiments will be reported in the figure captions. For all box plots throughout this work, boxes indicate the 25–75th percentiles, whiskers show the outlier values, and the midline indicates the median value. Reported *P* values were calculated using a two-tailed Student’s *t*-test for parametric distributions. Sample sizes are at least 3 independent experiments. No statistical test was used to determine sample sizes. The biological replicate (*n*) is defined as the number of independent experiments in which a new protein pattern was assembled. Independent experiments in some cases were performed on the same cover slip, which could fit up to six reaction chambers. Unless otherwise stated in the figure captions, the graphs show means ± s.d., and the error bars were calculated and are shown based on the number of independent experiments, as indicated. The distribution was assumed to be normal for all biological replicates.

